# The Molecular Basis of Monopolin Recruitment to the Kinetochore

**DOI:** 10.1101/456798

**Authors:** Rebecca Plowman, Namit Singh, Angel Payan, Eris Duro, Kevin D. Corbett, Adele L. Marston

## Abstract

In budding yeast meiosis I, the kinetochores of each sister chromatid pair are fused by the monopolin complex to mediate their monoorientation on the meiosis I spindle, enabling the biorientation and segregation of homologs. Monopolin forms a V-shaped complex with binding sites for the kinetochore protein Dsn1 at the apices of the V, suggesting that monopolin forms a physical bridge between the two sister kinetochores. Here, we reveal the molecular basis of the monopolin-kinetochore interaction and identify the key interfaces required for monopolin function at the kinetochore. The disordered N-terminus of budding-yeast Dsn1 unexpectedly possesses two binding motifs for the monopolin subunit Csm1, encompassing the previously-identified “Box 1” and “Box 2-3” regions of Dsn1. Strikingly, Dsn1 Box 1 and Box 2-3 bind the same conserved hydrophobic cavity on the monopolin complex subunit Csm1, suggesting that they are mutually exclusive for Csm1 binding, yet both regions are critical for monopolin function in *Saccharomyces cerevisiae* meiosis I. We find that Dsn1 Box 1 is an ancestral monopolin-binding motif that is conserved throughout fungi, including in the fission yeast *Schizosaccharomyces pombe*. In contrast, Box 2-3 is found only in species with sequence-defined point centromeres (*S. cerevisiae* and its close relatives), suggesting that this region contributes specifically to sister kinetochore crosslinking in meiosis I. Finally, we propose that phosphorylation of two conserved serine residues in Box 3 may stabilize monopolin at the kinetochore, providing a potential mechanism for enforcing specific sister kinetochore crosslinking in meiosis I.

## Introduction

Meiosis generates haploid gametes from a diploid progenitor cell through two consecutive rounds of chromosome segregation that follow a single round of DNA replication (reviewed in Duro and Marston (2015)). The first meiotic division (meiosis I) requires that the canonical chromosome segregation machinery be modified to direct the segregation of homologous chromosomes, rather than sister chromatids as in mitosis or meiosis II. Central to this process is the monoorientation of sister kinetochores, meaning that at metaphase I attachments are made to microtubules extending from the same spindle pole, rather than opposite poles, thereby ensuring the co-segregation of sister chromatids during anaphase I.

The mechanism of meiosis I sister kinetochore monoorientation is best understood in the budding yeast *Saccharomyces cerevisiae*. *S. cerevisiae* and its close relatives possess so-called “point centromeres,” compact sequence-defined centromeres that bind a single centromeric nucleosome and assemble a minimal kinetochore (Meraldi et al. 2006; Westermann et al. 2007; Gordon et al. 2011). In *S. cerevisiae* meiosis I, sister kinetochores are fused through the action of the kinetochore-binding monopolin complex, and together bind a single microtubule (Winey et al. 2005; Corbett et al. 2010; Corbett and Harrison 2012; Sarangapani et al. 2014). The conserved core of the monopolin complex comprises two nucleolar proteins, Csm1 and Lrs4 (Rabitsch et al. 2003). These proteins form a distinctive V-shaped complex, with two Csm1 homodimers bridged at their coiled-coil N-termini by a pair of Lrs4 subunits, thereby positioning two pairs of Csm1 globular-domain “heads” ~10 nm apart at the apices of the V (Corbett et al. 2010). Each Csm1 globular domain has a conserved hydrophobic cavity implicated in binding Dsn1, leading to the proposal that monopolin could bridge Dsn1 molecules from sister kinetochores to physically fuse the kinetochores (Corbett et al. 2010). Supporting this idea, kinetochore particles purified from cells in meiosis I bind microtubules more strongly than those from cells in mitosis or meiosis II, and this increased strength depends on the monopolin complex (Sarangapani et al. 2014). Further, addition of recombinant monopolin complex to kinetochores purified from mitotic cells increases their microtubule-attachment strength to match that of meiosis I kinetochores (Sarangapani et al. 2014).

A key unresolved question in monopolin function is how the complex specifically recognizes and crosslinks sister kinetochores. This specificity is likely mediated by two additional monopolin complex subunits, the meiosis-specific protein Mam1 and a CK1δ family kinase, Hrr25 (Toth et al. 2000; Rabitsch et al. 2003; Petronczki et al. 2006). Mam1, which is found only in point-centromere fungi, binds Csm1 and Hrr25 independently, through two flexibly-linked domains, thereby acting as a molecular tether to recruit Hrr25 to the monopolin complex (Corbett and Harrison 2012; Ye et al. 2016). While CK1δ family kinases are near-universal in eukaryotes, Hrr25 orthologs in point-centromere fungi possess a central domain that binds Mam1 and may uniquely regulate the protein’s kinase activity when it is associated with the monopolin complex (Ye et al. 2016). While the relevant substrates of monopolin-associated Hrr25 have not been identified, the flexibility and length (~120 Å) of the Mam1 tether would allow the kinase to access potential substrates within both monopolin and the kinetochore (Corbett and Harrison 2012; Ye et al. 2016). One candidate target is the kinetochore receptor for monopolin, Dsn1, which we previously showed is phosphorylated *in vitro* by Hrr25 (Ye et al. 2016). Hrr25’s kinase activity is dispensable for kinetochore localization of the monopolin complex *in vivo* (Petronczki et al. 2006) and for fusion of purified kinetochore particles *in vitro* (Sarangapani et al. 2014), but is required for sister kinetochore monoorientation in meiosis I (Petronczki et al. 2006). Together, these data suggest that kinetochore binding is functionally distinct from sister kinetochore crosslinking, and that Hrr25’s kinase activity is specifically important for the latter.

Apart from its critical role at meiosis I kinetochores, the Csm1-Lrs4 monopolin subcomplex acts as a molecular crosslinker in at least three other functional contexts in *S. cerevisiae*, some of which are likely conserved throughout fungi. Csm1 and Lrs4 reside in the nucleolus for the majority of the cell cycle, and a subset of Csm1-Lrs4 is released from the nucleolus after meiotic prophase to function at meiotic kinetochores (Rabitsch et al. 2003; Clyne et al. 2003). The complex is also released from the nucleolus in mitotic anaphase, when it localizes to kinetochores independently of Mam1 and Hrr25, and appears to suppress chromosome loss through an unknown mechanism (Brito et al. 2010). Within the nucleolus, Csm1 and Lrs4 are important for suppressing aberrant recombination within the highly-repetitive ribosomal DNA (rDNA) repeats, and are also required for Sir2-mediated transcriptional silencing of rDNA (Huang et al. 2006; Mekhail et al. 2008). Csm1 binds the nucleolar protein Tof2 through the same conserved hydrophobic cavity implicated in Dsn1 binding, and also binds a SUMO peptidase, Ulp2, in a structurally equivalent manner to Mam1 (Liang et al. 2017). Finally, we have recently identified another Csm1-binding protein, Dse3, which binds Csm1 equivalently to Mam1 and Ulp2 (Singh and Corbett 2018).The biological role of the Dse3-Csm1 interaction is not known.

Outside point-centromere fungi, Csm1 and Lrs4 are also important in chromosome and kinetochore organization and their molecular function is likely to be conserved. *S. pombe* Csm1 and Lrs4 (called Pcs1 and Mde4) prevent aberrant chromosome-microtubule attachments in mitosis (Gregan et al. 2007; Choi et al. 2009) and have been proposed to do so through either physical crosslinking of microtubule binding sites within a single kinetochore, or alternatively through recruitment of chromosome-organising condensin complexes to centromeric chromatin (Tada et al. 2011). Condensin-dependent organization of centromeres and rDNA is also thought to underlie the importance of Csm1-Lrs4 in the fungal pathogen *Candida albicans* (Burrack et al. 2013).

While the architecture of the monopolin complex and the structural basis for its interactions with numerous partners are known, direct molecular information about the monopolin-kinetochore interface is still lacking. Recent studies have identified a ~40-residue region within the disordered N-terminus of the core kinetochore protein, Dsn1, as the kinetochore receptor for the monopolin subunit Csm1 (Sarkar et al. 2013). This region, comprising residues 72-110 of *S. cerevisiae* Dsn1, is dispensable for vegetative growth but essential for sister kinetochore monoorientation in meiosis I (Sarkar et al. 2013). Sarkar et al. (2013) defined three conserved motifs in the Dsn1 72-110 region as Box 1, Box 2, and Box 3, and demonstrated their collective importance for Csm1 binding and monopolin function (Fig. 1a) (Sarkar et al. 2013). Here, we combine structural analysis of reconstituted Csm1-Dsn1 complexes with targeted mutagenesis, genetics, and imaging to dissect the molecular basis for monopolin recruitment and sister kinetochore monoorientation. We find that the Dsn1 Box 1 and Box 2-3 regions can each bind the conserved hydrophobic cavity on Csm1, and that these two interaction modes are mutually exclusive in a given Csm1-Dsn1 complex. We demonstrate that both interfaces are required for robust monopolin recruitment to kinetochores and sister kinetochore monoorientation, and that simultaneous disruption of both interfaces leads to additive effects on meiosis. Finally, we show that Dsn1 Box 1 is widely conserved in fungi and provide evidence, using *S. pombe* proteins, that Box1 is the ancestral kinetochore receptor for monopolin. The Dsn1 Box 2-3 region, meanwhile, is conserved only in point-centromere fungi and likely represents an adaptation to the complex’s meiotic functions. Further, Dsn1 Box 3 contains two conserved serine residues that are likely to be phosphorylated to modulate Dsn1-Csm1 binding, providing a potential molecular mechanism for sister kinetochore crosslinking specificity in meiosis I.

**Fig. 1.**
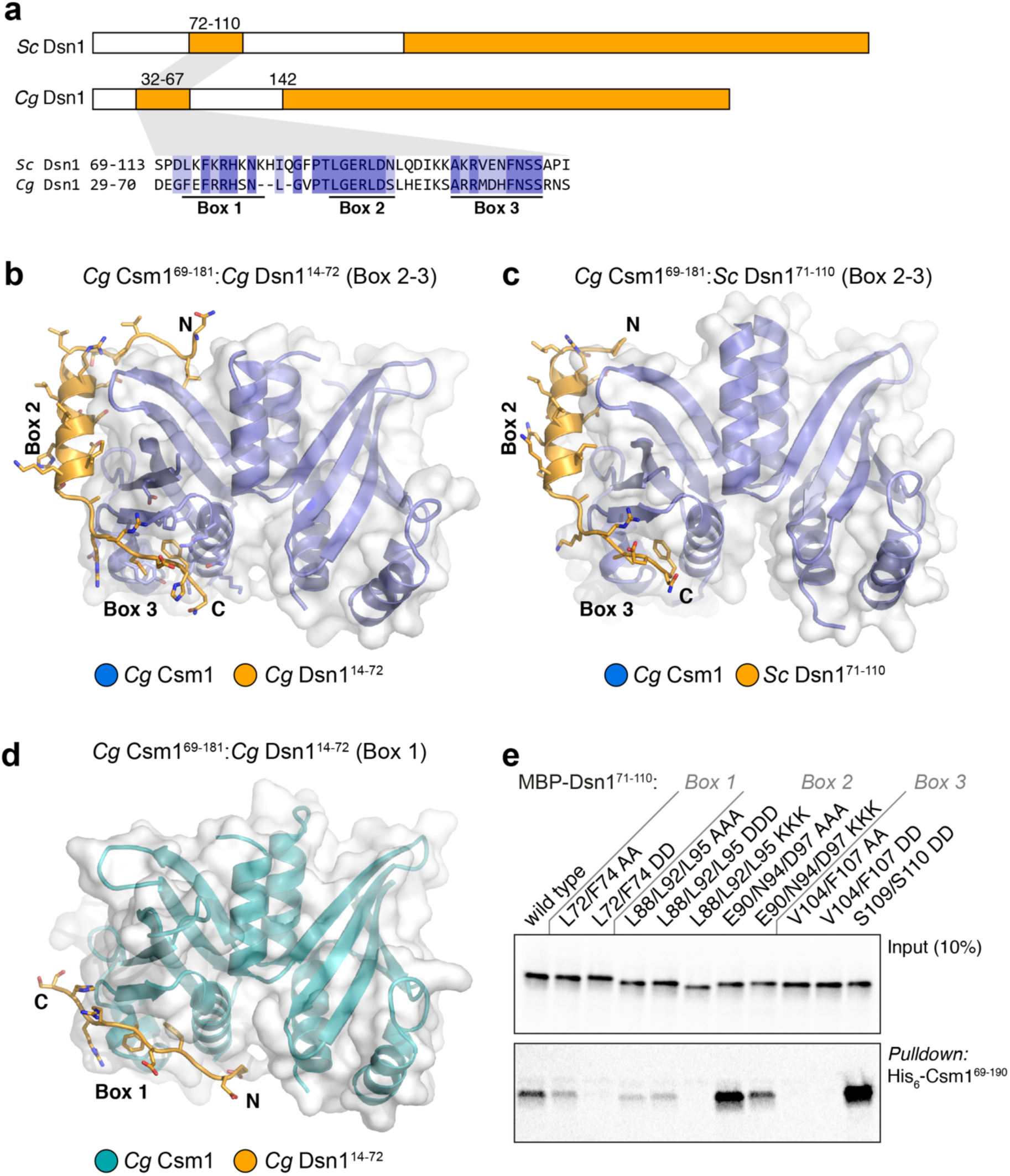
Structure of the Csm1-Dsn1 complex. **a** Domain schematic of Dsn1 from *S. cerevisiae* and *C. glabrata*, with conserved regions shown in orange. The Dsn1 C-terminal domain forms a folded complex with other MIND complex subunits, while the N-terminal conserved region interacts with Csm1. *Bottom:* Sequence alignment of the *S. cerevisiae* and *C. glabrata* Box 1-2-3 region; **b** Overall view of the *Cg* Csm1^69-181^:*Cg* Dsn1^14-72^ complex, showing the Dsn1 Box 2-3 region (orange) interacting with a Csm1 dimer (blue with white surface); **c** Overall view of the *Cg* Csm1^69-181^:*Sc* Dsn1^71-110^ complex, showing the Dsn1 Box 2-3 region. See Fig. S2 for more details on Csm1-Dsn1 Box 2-3 interactions, and Fig. S3 for crystal packing interactions; **d** Overall view of the *Cg* Csm1^69-181^:*Cg* Dsn1^14-72^ complex, showing the Dsn1 Box 1 region (orange) interacting with a Csm1 dimer (teal with white surface). See Fig. S2a for crystal packing interactions for this complex; **e** Ni^2+^-pulldown of in vitro translated *S. cerevisiae* Dsn1 N-terminal region constructs by *Sc* His_6_-Csm1^69-190^

## Materials and Methods

### Cloning and Protein Purification

All protein coding sequences were amplified from genomic DNA and cloned into pET-based vectors, either without tags or encoding N-terminal TEV protease-cleavable His_6_ or His_6_-SUMO tags. Coexpression cassettes were generated by PCR and re-inserted into the same vectors. Point-mutations were generated by PCR. For expression, vectors were transformed into *E. coli* Rosetta2 (DE3) pLysS cells (EMD Millipore), and cultures were grown at 37°C to an absorbance at 600 nm of ~0.8. The cultures were shifted to 20°C and protein expression was induced by the addition of 0.25 mM IPTG, and cells were grown ~16 hours before harvesting by centrifugation. For protein purification, cells were resuspended in protein buffer (20 mM Tris-HCl pH 7.5, 5% glycerol, 2 mM β-mercaptoethanol) plus 300 mM NaCl and 10 mM imidazole, lysed by sonication, and centrifuged 30 min at 17,000 RPM to remove cell debris. The supernatant was loaded onto a 5 mL Histrap HP column (GE Life Sciences), washed with protein buffer plus 300 mM NaCl/20 mM imidazole, then with protein buffer plus 100 mM NaCl/20 mM Imidazole. Protein was eluted with protein buffer plus 100 mM NaCl/250 mM imidazole. Protein was then loaded onto a 5 mL Hitrap Q HP column (GE Life Sciences), washed with protein buffer plus 100 mM NaCl, then eluted with a gradient to 600 mM NaCl. Peak fractions were pooled, and TEV protease (Tropea et al. 2009) was added to cleave His_6_ or His_6_-SUMO tags, and the mixture was incubated 16 hours at 4°C (for *Cg*Csm1^69-181^:*Sc* His_6_-Dsn1^71-110^ and *Cg* His_6_-Csm1^69-181^:*Sc*Dsn1^71-110^, tag cleavage was not performed; eluted fractions were instead concentrated and passed directly over a Superdex 200 column). After tag cleavage, the mixture was passed over Histrap HP and the flow-through collected, concentrated by ultrafiltration (Amicon Ultra, EMD Millipore), then passed over a HiLoad Superdex 200 column size exclusion column (GE Life Sciences) in protein buffer plus 300 mM NaCl (with 1 mM dithiothreitol substituting for β-mercaptoethanol) for final purification. Protein was concentrated to ~10 mg/mL and stored at 4°C for crystallization.

### Crystallization and Structure Determination

#### *Cg*Csm1^69-181^: *Cg*Mam1^162-216^

For crystallization of the *Cg*Csm1^69-181^:*Cg*Mam1^162-216^ complex, purified protein at 10 mg/mL was mixed 1:1 with well solution containing 0.1 M MES pH 6.5, 0.6 M NaCl, and 20% PEG 4000. Crystals were cryoprotected with the addition of 20% PEG 400 and flash-frozen in liquid nitrogen. Diffraction data were collected to 3.03 Å resolution at the Advanced Photon Source, NE-CAT beamline 24ID-E (support statement below) and indexed/reduced with the RAPD automated data-processing pipeline (https://github.com/RAPD/RAPD), which uses XDS (Kabsch 2010) for indexing and integration, and the CCP4 programs AIMLESS (Evans and Murshudov 2013) and TRUNCATE (Winn et al. 2011) for scaling and structure-factor calculation. The structure was determined by molecular replacement in PHASER (McCoy et al. 2007) using the structure of *S. cerevisiae* Csm1 (PDB ID 3N4R) (Corbett et al. 2010) as a search model. The model, including all *Cg*Mam1 residues, was manually built in COOT (Emsley et al. 2010) and refined in phenix.refine (Afonine et al. 2012) using positional, individual B-factor, and TLS refinement (Table S1).

#### *Cg*Csm1^69-181^: *Cg*Dsn1^14-72^

For crystallization of the *Cg*Csm1^69-181^:*Cg*Dsn1^14-72^ complex, purified protein at 10 mg/mL was mixed 1:1 with well solution containing 0.45 M Ammonium sulfate, 5% PEG 3350, and 0.1 M Bis-Tris, pH 5.5 in hanging-drop format at 20°C. Crystals were cryoprotected with the addition of 25% glycerol and flash-frozen in liquid nitrogen. Diffraction data were collected to 2.27 Å (native) or 2.59 Å (selenomethionine) resolution at the Advanced Photon Source, NE-CAT beamline 24ID-E and indexed/reduced with the RAPD automated data-processing pipeline. The structure was determined by molecular replacement, manually rebuilt and refined as above.

#### *Cg* His_6_-Csm1^69-181^:*Sc*Dsn1^71-110^

For crystallization of the *Cg* His_6_-Csm1^69-181^:*Sc*Dsn1^71-110^ complex (form 2), purified protein at 10 mg/mL was mixed 1:1 with well solution containing 0.2 M MgCl_2_, 0.1 M Tris-HCl pH8.5, and 25% PEG 3350. Crystals were cryoprotected with the addition of 20% PEG 400 and flash-frozen in liquid nitrogen. Diffraction data were collected to 2.5 Å resolution at the Stanford Synchrotron Radiation Laboratory, beamline 14-1 (support statement below). Data were indexed, reduced, and scaled with HKL2000 (Otwinowski and Minor 1997) and converted to structure factors using TRUNCATE (Winn et al. 2011). The structure was determined by molecular replacement, manually rebuilt and refined as above.

#### *Cg*Csm1^69-181^: *Cg*Dsn1^43-67^DD

For crystallization of the *Cg*Csm1^69-181^:*Cg*Dsn1^43-67^DD complex (serines 66 and 67 mutated to aspartate), purified protein at 10 mg/mL was mixed 1:1 with well solution containing 0.1M Sodium acetate pH 4.5 and 3 M NaCl. Crystals were cryoprotected with the addition of 2.5 M Sodium Malonate pH 4.5 and flash-frozen in liquid nitrogen. Diffraction data were collected were collected at the Advanced Photon Source, NE-CAT beamline 24ID-E and indexed/reduced with the RAPD automated data-processing pipeline. The structure was determined by molecular replacement, manually rebuilt and refined as above.

All macromolecular structure figures were generated with PyMOL version 2.2 (Schrödinger, LLC), and surface charge for Fig. 3e was calculated using the APBS (Jurrus et al. 2018) plugin for PyMOL.

### Synchrotron Support Statements

#### Advanced Photon Source

This work is based upon research conducted at the Northeastern Collaborative Access Team beamlines, which are funded by the National Institute of General Medical Sciences from the National Institutes of Health (P30 GM124165). The Eiger 16M detector on 24-ID-E beam line is funded by a NIHORIP HEI grant (S10OD021527). This research used resources of the Advanced Photon Source, a U.S. Department of Energy (DOE) Office of Science User Facility operated for the DOE Office of Science by Argonne National Laboratory under Contract No. DE-AC02-06CH11357.

#### Stanford Synchrotron Radiation Lightsource

Use of the Stanford Synchrotron Radiation Lightsource, SLAC National Accelerator Laboratory, is supported by the U.S. Department of Energy, Office of Science, Office of Basic Energy Sciences under Contract No. DE-AC02-76SF00515. The SSRL Structural Molecular Biology Program is supported by the DOE Office of Biological and Environmental Research, and by the National Institutes of Health, National Institute of General Medical Sciences (including P41GM103393). The contents of this publication are solely the responsibility of the authors and do not necessarily represent the official views of NIGMS or NIH.

### Protein-protein interaction assays

For in vitro translation and Ni^2+^ pulldown assays, *S. pombe* Mis13 (Dsn1) residues 1-100 and *S. cerevisiae* Dsn1 residues 71-110 (and point mutants thereof) were cloned with an N-terminal maltose binding protein tag (no His_6_-tag) into a pET-based vector with a Kozak sequence immediately upstream of the coding sequence. These vectors were used as a template for in vitro transcription/translation using a TNT T7 coupled transcription/translation kit (Promega) in the presence of ^35^S-labeled methionine to generate prey proteins for pulldowns. Ten µl of transcribed protein mix was incubated with 10 µg His_6_-tagged bait protein (*S. pombe* Csm1^125-261^ or *S. cerevisiae* Csm1^69-190^) in 50 µl buffer (20 mM HEPES, pH 7.5, 150 mM NaCl, 20 mM imidazole, 5% glycerol, 1 mM dithiothreitol (DTT), 0.1% NP-40) for 90 min at 4°C, then 15 µl Ni-NTA beads were added, and the mixture was incubated a further 45 min. Beads were washed three times with 0.5 ml buffer, then eluted with 25 ml elution buffer (2x SDS-PAGE loading dye plus 400 mM imidazole) and boiled. Samples were run on SDS-PAGE, dried, and scanned with a phosphorimager. For fluorescence polarization peptide-binding assays, purified *S. pombe* Csm1^125-261^ (wild-type or I241D mutant, equivalent to *Sc* Csm1 I161D) at 20 nM-250 µM was incubated with 20 nM *Sp* Mis13 5-17 peptide (fluorescein isothiocyanate-labelled at its N-terminus) in a buffer containing 20 mM Tris 7.5, 300 mM NaCl, 10% glycerol, 0.01% NP-40, and 1 mM DTT (50 µL reactions, measured in triplicate). Binding data were fit to a single-site binding model with Prism version 7 (Graphpad Software).

### Isothermal Titration Calorimetry

Isothermal titration calorimetry was performed on a Microcal ITC 200 (Malvern Panalytical) in protein buffer plus 300 mM NaCl and 1 mM dithiothreitol. His_6_-MBP-fused Dsn1 fragments at 1-1.5 mM were injected into a sample cell containing untagged Csm1 at 100-200 µM.

### Yeast strains and plasmids

Yeast strains used in this study were derivatives of SK1 and are given in Table S2. The *CEN5*-GFP marker consists of two components: (1) an array of *tet* operator sequences inserted at the chromosome V centromere, and (2) a Tet repressor protein fused to GFP, which binds to and specifically marks these operator sites and were previously described in (Toth et al. 2000). *MAM1-9MYC* was also described in (Toth et al. 2000). *PDS1-tdTomato* and *pCLB2-CDC20* were described in (Lee and Amon 2003) and (Matos et al. 2008), respectively. *MTW1-tdTomato* was generated in SK1 as described in (Fernius et al. 2013). Point mutations in *DSN1-6His-3FLAG* were generated in plasmid pSB1590 (Akiyoshi et al. 2010) using the Quick Change II-XL kit (Agilent Technologies) and integrated into the *DSN1* endogenous locus by PCR-mediated transformation. Plasmids generated in this study are given in Table S3.

### Yeast growth conditions

Sporulation was induced as described by (Vincenten et al. 2015). Briefly, diploid yeast were grown overnight on YPG agar (1% yeast extract, 2% Bacto peptone, 2.5% glycerol, and 2% agar), transferred to YPD4% agar (1% yeast extract, 2% Bacto peptone, 4% glucose, and 2% agar) and incubated for 24 h before inoculating into YEPD liquid medium (1% yeast extract, 2% Bacto peptone, and 2% glucose) and incubating with shaking for 24 h. Cells were transferred to BYTA (1% yeast extract, 2% Bacto tryptone, 1% potassium acetate, 50mM potassium phthalate) at an OD_600_ = 0.2 – 0.3 or YPA (1% Yeast Extract, 2% Tryptone peptone, 1% potassium acetate) and incubated for a further ~16 h. Cells were washed once with sterile distilled water and re-suspended in SPO medium (0.3% potassium acetate, pH 7) at an OD_600_ = 1.8 – 1.9; t=0. Cells were incubated at 30°C for the duration.

### Chromatin immunoprecipitation qPCR

Cells carrying *pCLB2-CDC20* (Lee and Amon 2003) were induced to sporulate. After 6 h in SPO, cells were then fixed in 1% formaldehyde for 1 h, washed twice with TBS (20 mM Tris-HCl pH7.5, 150 mM NaCl) and once with 1x FA lysis buffer (50mM HEPES-KOH at pH 7.5, 150mM NaCl, 1mM EDTA, 1% v/v Triton X-100, 0.1% w/v Sodium Deoxycholate) containing 0.1% w/v SDS before resuspending in 1x FA lysis buffer/0.1% SDS. Cells were lysed in a Fastprep Bio-pulverizer FP120 with silica beads (Biospec Products). Samples were sonicated to fragment chromosomal DNA using a BioRupter (Diagenode). Aliquots of the resultant chromatin solution were incubated with either anti-Myc (9E10, Biolegend) or anti-FLAG (M2, Sigma) antibodies and Protein G Dynabeads (Life Technologies) overnight at 4°C. Following sequential washes with CB1 (FA lysis buffer/0.1% SDS/ 275 mM NaCl), CB2 (FA lysis buffer/0.1% SDS/ 500 mM NaCl), CB3 (10 mM Tris-HCl, pH 8, 0.25 M LiCl, 1mM EDTA, 0.5% NP-40, 0.5% Na Deoxycholate) and CB4 (TE: 10 mM Tris-HCl, pH 8, 1 mM EDTA), immunoprecipitates and 1/100 input chromatin were recovered by boiling (10 min) with a 10% slurry of Chelex-100 resin before adding Proteinase K (0.125 mg) and incubating at 55°C for 30 min, then boiled for a further 10 min. Samples were centrifuged and supernatant taken for qPCR. qPCR was performed on a on a Roche Lightcycler with LUNA universal qPCR Master Mix (New England Biolabs). Primers used for qPCR are given in Table S4. To calculate ChIP enrichment/input, ΔCT was calculated according to: ΔCT = (CT_(ChIP)_ − [CT_(Input)_− logE (Input dilution factor)]) where E represents the specific primer efficiency value. Enrichment/ input value was obtained from the following formula: E^^−ΔCT^. qPCR was performed in triplicate from three or more independent cultures. Error bars represent standard error. Figures show the mean values for each strain, averaged over all individual experiments and biological replicates. Wild type and no tag controls were included for reference in all individual experiments and replicates. The number of replicates for each strain is indicated in the figure legends.

### Western Blotting

Samples for immunoblot analysis were fixed in 5% TCA and cell pellets were washed once with acetone. Cells were lysed in 50 mM Tris (pH 7.5), 1 mM EDTA, and 50 mM DTT containing protease inhibitors with glass beads, boiled in 1× sample buffer and visualized by detection of chemiluminesence on autoradiograms. Mouse Anti-FLAG M2 antibodies (Sigma) and mouse Anti-cMYC (9E10, Biolegend) were used at 1:1000 dilution, and rabbit Anti-PGK1 (Marston lab stock) was used at 1:10,000 dilution.

### Spore viability

Haploid yeast strains with the relevant genotypes were mated and single diploid colonies were incubated on SPO agar. A minimum of two diploid isolates were chosen for spore dissection. The total number of tetrads dissected for each strain is indicated in the figure legend. Spores were allowed to grow for 2 days on YPDA at 30°C before scoring the number of viable colonies per tetrad.

### Live cell imaging

Cells were induced to sporulate as above. Following 2 h incubation in SPO medium in flasks, cells were immobilized on Concanavalin A-coated cover slips in ibidi 4-well or 8-well dishes, fresh sporulation media was added to the dish and imaging commenced. Imaging was performed on a Zeiss Axio Observer Z1 (Zeiss UK, Cambridge) equipped with a Hamamatsu Flash 4 sCMOS camera, Prior motorised stage and Zen 2.3 acquisition software. Images were processed in image J and 8 Z-stacks were projected to maximum intensity.

## Results

### Reconstitution and structure of a budding-yeast Csm1-Dsn1 complex

To better understand the interactions between the monopolin complex and the kinetochore, and the roles of the Dsn1 Box 1, 2, and 3 regions in Csm1 binding, we sought to reconstitute a complex between Csm1 and the Dsn1 N-terminus. We first separately purified the *S. cerevisiae* Csm1 globular domain (residues 69-190 of 190) and the Dsn1 Box 1-2-3 region (residues 71-110 of 534) and measured a binding affinity (*K_d_*) of 12 µM by isothermal titration calorimetry (ITC) (Fig. S1a). We next co-expressed and purified a stable *S. cerevisiae* (*Sc*) Csm1^69-181^:Dsn1^71-110^ complex (with the C-terminal 9 disordered residues of Csm1 removed), but were unable to identify crystallization conditions for this complex. We therefore screened paralogs from several related budding yeast, and successfully purified a complex between the *Candida glabrata* (*Cg*) Csm1 globular domain (residues 69-181) and the Dsn1 Box 1-2-3 region (residues 14-72) (Fig. S1b, c). We identified crystallization conditions for this complex and determined the structure to 2.3 Å resolution (Table S1). The *Cg* Dsn1 Box 1-2-3 region shows high homology with the equivalent region of *Sc* Dsn1 (56% identity and 82% similarity between *Sc* Dsn1 residues 72-110 and *Cg* Dsn1 residues 32-67) (Fig. 1a), and we were able to reconstitute a complex of *Cg* Csm1^69-181^ (56% identical to *Sc* Csm1 in this region) with *Sc* Dsn1^71-110^ (Fig. S1d). We crystallized and determined the structure of this chimeric complex to 2.5 Å resolution. We also determined a 3.0 Å-resolution structure of *Cg* Csm1^69-181^ in complex with the Csm1-binding region of *Cg* Mam1 (residues 162-216) (Fig. S1e). While our attempts to purify a ternary complex of Csm1, Dsn1, and Mam1 were unsuccessful, these crystal structures provide a comprehensive picture of how budding yeast Csm1 interacts through its C-terminal globular domain with Dsn1 and Mam1.

Our prior biochemical data showed that *Sc* Dsn1 interacts with a highly conserved hydrophobic cavity on the Csm1 globular domain (Corbett et al. 2010). Later work implicated the Dsn1 Box 1-2-3 region as necessary for binding Csm1, and mutagenesis revealed a particular requirement for the Box 2-3 region (Sarkar et al. 2013). Our structures of the *Cg* Csm1^69-181^:Dsn1^14-72^ (Fig. 1b) and *Cg* Csm1^69-181^:*Sc* Dsn1^71-110^ (Fig. 1c) complexes reveal a consistent interface between Dsn1 Box 2-3 and Csm1 (Fig. S2), while the *Cg* Csm1^69-181^:Dsn1^14-72^ structure (Fig. 1c) reveals a second interface between Csm1 and Dsn1 Box 1 (Fig. 1d, S3a). Therefore, all three conserved segments in the Dsn1 N terminus contact Csm1. Intriguingly, the conserved hydrophobic cavity on Csm1 was involved in binding both Dsn1 Box 3 (in both the *Cg* Csm1^69-181^:Dsn1^14-72^ and *Cg* Csm1^69-181^:*Sc* Dsn1^71-110^ structures) and Box 1 (in the *Cg* Csm1^69-181^:Dsn1^14-72^ structure), which form strikingly similar interfaces with Csm1 (Fig. 1b-d). We next sought to understand which of these interfaces are important for sister kinetochore monoorientation during meiosis.

### Dsn1 Box 2 contributes to successful meiosis

In the crystal structures of both *Cg* Csm1^69-181^:Dsn1^14-72^ and the chimeric *Cg* Csm1^69-181^:*Sc* Dsn1^71-110^ complex, the Dsn1 Box 2-3 region wraps around the Csm1 globular domain, with Box 2 forming an α-helix that packs against the “side” of the Csm1 dimer, and Box 3 binding the Csm1 hydrophobic cavity (Fig. 1b, c; Fig. S2). Box 2 is highly conserved in yeast with point centromeres, with an alternating pattern of hydrophobic (*Sc* Dsn1 L88/L92/L95) and polar (*Sc* Dsn1 E90/N94/D97) residues (Fig. 2a). In both structures, this region forms an α-helix oriented with the hydrophobic residues facing outward into solution, and the polar residues packed tightly against Csm1 (Fig. 2a). This binding mode is unexpected, as hydrophobic residues are most often buried in protein-protein interfaces, rather than solvent-exposed. To determine the importance of Dsn1 Box 2 for Csm1 binding and successful meiosis, we mutated either the polar or hydrophobic residues in Box 2 and tested their function *in vivo* and *in vitro*. First, we produced the Dsn1 Box 1-2-3 region (71-110) by *in vitro* translation, and performed pulldown assays with purified Csm1 (Fig. 1e). This assay showed that mutation of the polar residues contacting Csm1 (Dsn1 E90, N94, and D97) to lysine did not detectably reduce Csm1 binding, while mutation to alanine appeared to increase binding (Fig. 1e). In contrast, mutation of the solvent-exposed hydrophobic residues of Dsn1 Box 2 (L88, L92, and L95) impaired binding to Csm1, with lysine substitutions having the greatest effect and alanine or aspartate substitutions causing a modest reduction in binding (Fig. 1e). This suggests that, at least when Dsn1 Box 1 and 3 are present, mutations in the Csm1-contacting surface of Box 2 do not compromise binding, while the solvent-exposed hydrophobic residues play an unexpectedly important role in Csm1 binding.

**Fig. 2.**
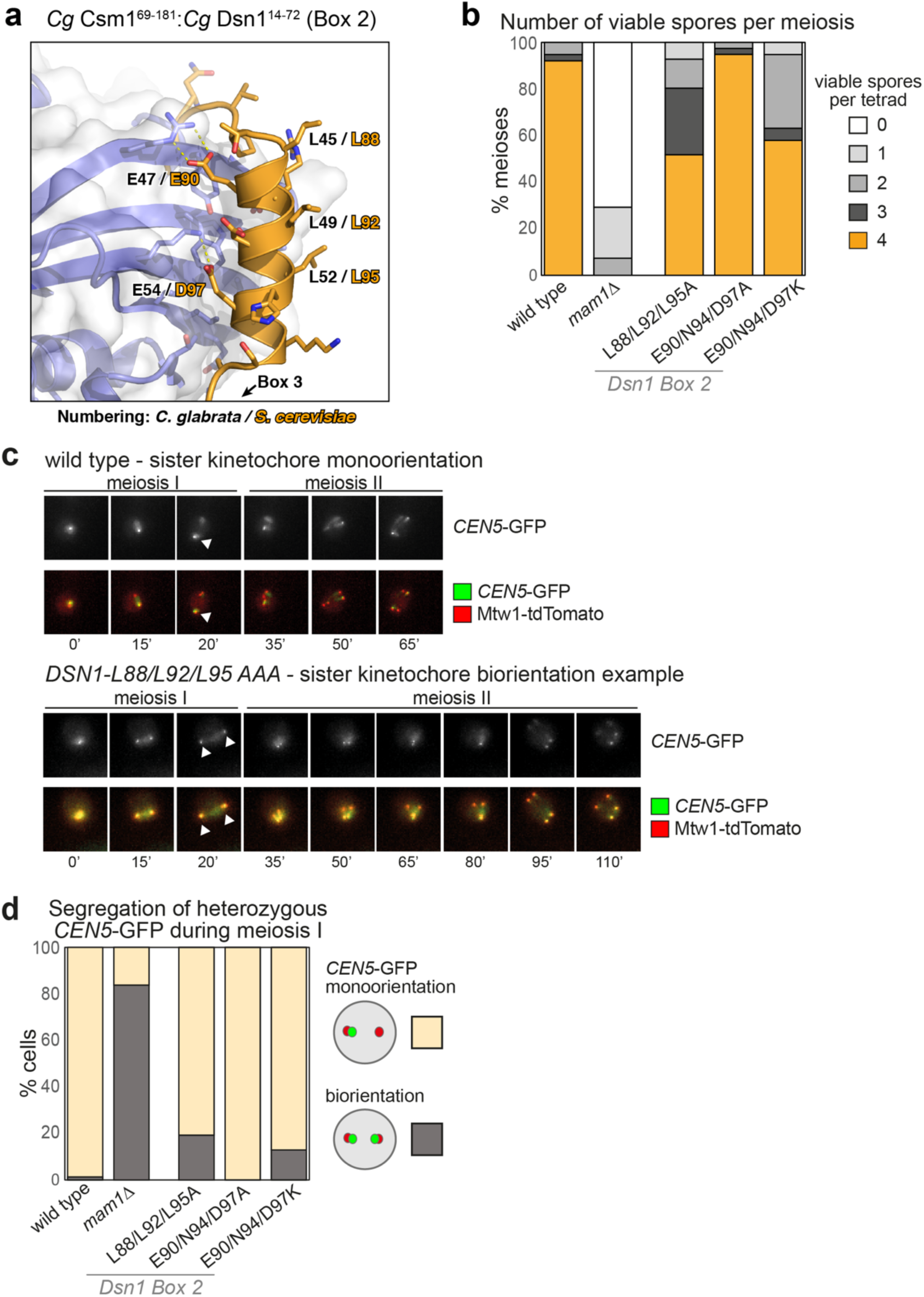
Dsn1 Box2 contributes to successful meiosis. **a** Closeup view of the *Cg* Dsn1 Box 2 region (orange) interacting with the “side” of a Csm1 protomer (blue with white surface) in the *Cg* Csm1^69-181^:*Cg* Dsn1^14-72^ complex. Residue numbers shown are for *Cg* Dsn1, with *Sc* Dsn1 equivalents shown in orange text. See Fig. S2d-f for equivalent views of the *Cg* Csm1^69-181^:*Sc* Dsn1^71-110^ and *Cg* Csm1^69-181^:*Cg* Dsn1^43-67^DD complexes; **b** Point mutations in Dsn1 affect spore survival. Diploid cells carrying the indicated homozygous mutations in *DSN1* were sporulated, dissected, and the number of spores that formed colonies from each tetrad was scored. Between 38 and 56 tetrads were dissected for each condition, from a minimum of two independent diploid strains. Diploid strains used were generated from matings between AMy1827 and AMy1828 or AMy1835 (wild type), AMy1932 and AMy1947 (*mam1**Δ***), AMy21921 and AMy22719 (*DSN1-L88A L92A L95A*), AMy23151 and AMy23152 (*DSN1-E90A N94A D97A*), and AMy24629 and AMy24632 (*DSN1-E90K N94K D97K*); **c, d** Live cell imaging of heterozygous *CEN5-*GFP foci during meiosis reveals defective monoorientation in the presence of Dsn1 Box 2 mutations. Cells also carry Mtw1-tdTomato to label kinetochores and Pds1-tdTomato, the destruction of which marks anaphase I onset; **c** Representative images of strains producing either wildtype Dsn1 or Dsn1-L88A L92A L95A. While wild type cells segregate a single *CEN5*-GFP focus to one pole, some *DSN1-L88A L92A L95A* cells split GFP foci and exhibit delayed meiosis II. Arrowheads indicate position of *CEN5-GFP* foci during anaphase I, revealing whether they segregate to the same pole (monooriented, as in the wild type example) or opposite poles (bioriented as in *DSN1-L88A L92A L95A* cells). Images are from frames taken at 15 min intervals; **d** Scoring of GFP foci position at anaphase I onset, defined as the first occasion on which Mtw1-tdTomato segregate. Strains used were AMy25832 (wild type; n= 78), AMy25881 (*DSN1-L88A L92A L95A*; n=26), AMy25763 (*DSN1-E90A N94A D97A*; n=39) and AMy25881 (*DSN1-E90K N94K D97K*; n=93)

To determine the role of Dsn1 Box 2 in meiosis, we generated *S. cerevisiae* strains with mutations in either the Csm1-contacting polar residues (Dsn1 E90A/N94A/D97A and Dsn1 E90K/N94K/D97K) or the solvent-exposed hydrophobic residues (Dsn1 L88A/L92A/L95A). Impaired monopolin function causes mis-segregation of chromosomes during meiosis, producing aneuploid gametes which are frequently inviable. Therefore, we first analysed the ability of *S. cerevisiae* strains with homozygous mutations in Dsn1 to produce viable meiotic progeny, or spores. Consistent with the Csm1 binding studies, we observed that mutation of the Dsn1 Box 2 polar residues to alanine (Dsn1-E90A/N94A/D97A) had no detectable effect on spore viability, while lysine substitutions (Dsn1-E90K/N94K/D97K) resulted in reduced spore viability (Fig. 2b). Mutation of the solvent-exposed hydrophobic residues (Dsn1-L88A/L92A/L95A), which strongly affected Csm1 binding *in vitro*, also reduced spore viability (Fig. 2b). We next asked whether the ability to establish sister kinetochore monoorientation during meiosis I could underlie these effects on spore viability. We imaged live *DSN1-L88A/L92A/L95A, DSN1-E90A/N94A/D97A*, and *DSN1-E90K/N94K/D97K* cells carrying a heterozygous *CEN5-*GFP marker (which tracks the segregation of a single sister chromatid pair in meiosis I; see Methods), a kinetochore marker (Mtw1-tdTomato), and a marker for anaphase I onset (Pds1-tdTomato). Using these strains, we detected monoorientation defects consistent with each mutation’s effect on spore viability: While virtually all wild type and *DSN1-E90A/N94A/D97A* cells segregate *CEN5-*GFP foci to the same pole during anaphase I, segregation of *CEN5-*GFP foci to opposite poles was observed for ~20% of *DSN1-L88A/L92A/L95A* and *E90K/N94K/D97K* cells during anaphase I (Fig. 2c, d). While these effects are less severe than observed in a *mam1Δ* mutant (Fig. 2d), the data nonetheless indicates that Dsn1 Box 2 is important for co-segregation of sister chromatids during anaphase I.

Although we cannot rule out that the amino acid changes in Dsn1-L88A/L92A/L95A cause structural perturbations of the Dsn1 Box2 α-helix, these findings suggest, unexpectedly, that the hydrophobic outer surface is critical for sister chromatid co-segregation during meiosis I. Interestingly, the polar residues on the Csm1-binding inner surface of the Dsn1 Box 2 α-helix can be mutated to alanine without affecting spore viability or co-segregation of sister kinetochores, at least in the presence of functional Box 1 and 3.

### Dsn1 Box 3 is critical for meiosis

While Dsn1 Box 2 forms an α-helix and associates with the “side” of the Csm1 dimer, Box 3 forms an extended conformation that packs tightly against the Csm1 conserved hydrophobic cavity (Fig. 3a; Fig. S2a-c, g-h). This binding is equivalent to the nucleolar protein Tof2, which we previously showed shares limited sequence homology with Dsn1 Box 3 (Liang et al. 2017) (Fig. S4). The core of the interaction comprises two conserved hydrophobic residues (*Sc* Dsn1 V104 and F107) inserted into the conserved hydrophobic cavity on Csm1. These residues are bracketed by positively charged amino acids (*Sc* Dsn1 K102 and R103) on the N-terminal side, and highly-conserved serine residues (*Sc* Dsn1 S109 and S110) on the C-terminal side (Fig. 3a; Fig. S3g-h). Mutations in the hydrophobic residues either to alanine (*DSN1-V104A/F107A*) or aspartate (*DSN1-V104D/F107D*) abolished Csm1 binding *in vitro* (Fig. 1e). Consistently, Dsn1 Box 3 mutations V104A/F107A and V104D/F104D led to a marked decrease in spore viability, whether present in single copy (heterozygous) or both copies (homozygous) (Fig. 3b). We also observed increased separation of *CENV-*GFP labeled sister chromatids to opposite poles in anaphase I with these mutants (Fig. 3c). Finally, we measured monopolin complex recruitment to kinetochores *in vivo* by analyzing Mam1 association with a representative kinetochore by chromatin immunoprecipitation (ChIP). This assay revealed that homozygous Dsn1 Box 3 mutations (either *DSN1-V104A/F107A* or *DSN1-V104D/F107D*) caused a significant reduction in monopolin complex association with kinetochores compared to wild type cells. Importantly, these effects were not due to defective kinetochore assembly, as Dsn1 Box 3 mutations did not affect overall Dsn1 levels (Fig. S5a), and kinetochore association of both Dsn1 itself and the KMN-network protein Ndc80 was unaffected (Fig. S5b, c). Collectively, these findings establish that the interface between Dsn1 Box 3 and Csm1 is critical for sister kinetochore co-segregation during meiosis I.

**Fig. 3.**
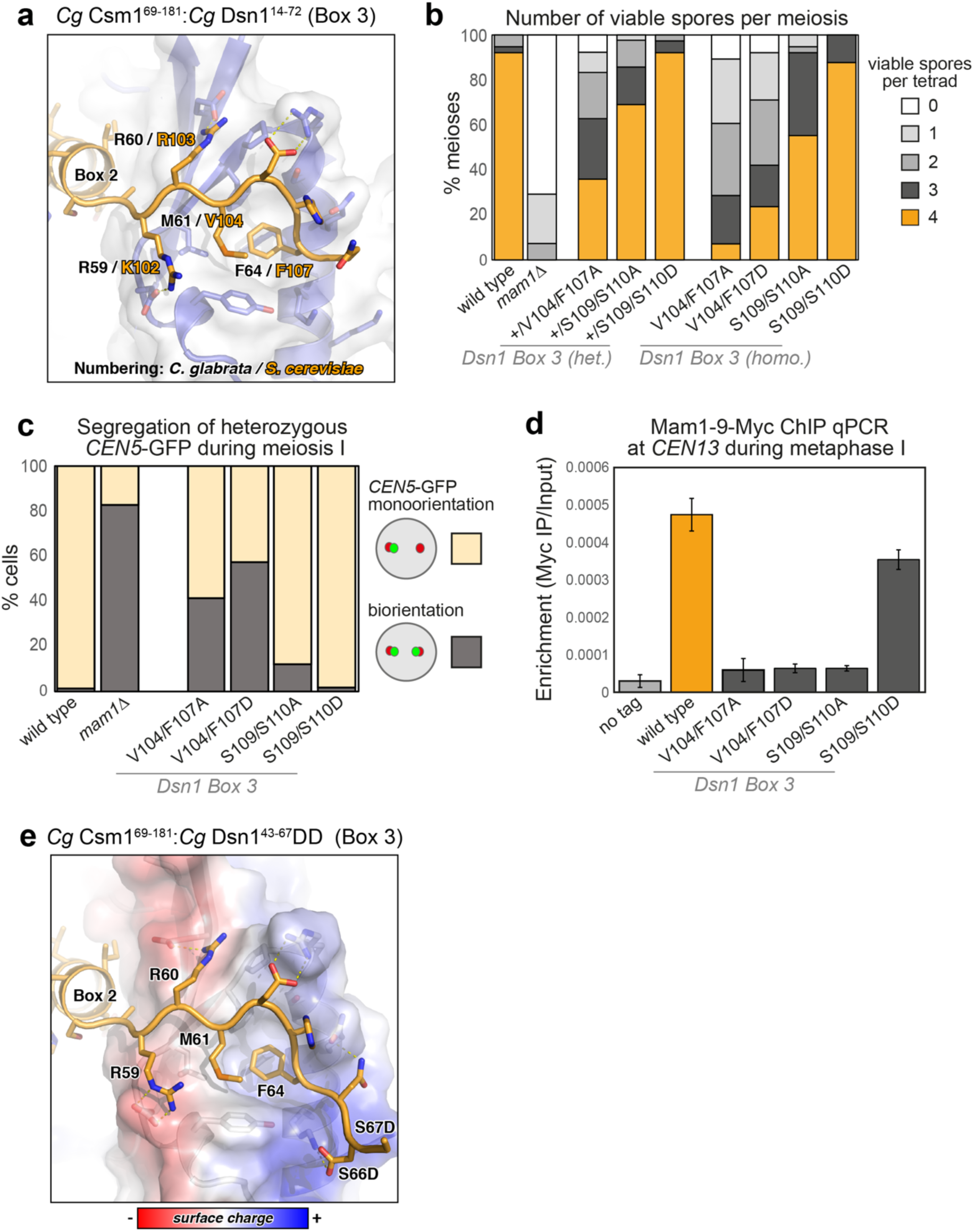
Dsn1 Box 3 residues are critical for meiosis. **a** Closeup view of the *Cg* Dsn1 Box 3 region (orange) interacting with the Csm1 conserved hydrophobic cavity (blue with white surface) in the *Cg* Csm1^69-181^:*Cg* Dsn1^14-72^ complex. Residue numbers shown are for *Cg* Dsn1, with *Sc* Dsn1 equivalents shown in orange text. See Fig. S2g-i for equivalent views of the *Cg* Csm1^69-181^:*Sc* Dsn1^71-110^ and *Cg* Csm1^69-181^:*Cg* Dsn1^43-67^DD complexes; **b** Diploid cells with heterozygous or homozygous mutations in *DSN1* were sporulated, dissected and the number of spores which grew up from each tetrad scored. Between 38 and 78 tetrads were dissected for each condition, from a minimum of two independent diploids. Data for wild type and *mam1Δ* is reproduced from Fig. 2b. Heterozygous diploids were generated from crosses between AMy1827 and AMy24652 (*DSN1-V104F F107A*), AMy1827 and AMy26803 (*DSN1-S109A S110A*), AMy1827 and AMy24744 (*DSN1-S109D S110D*). Homozygous diploids were generated from crosses between AMy24624 and AMy24652 (*DSN1-V104F F107A*), AMy26426 and AMy26803 (*DSN1-S109A S110A*), AMy24744 and AMy24688 (*DSN1-S109D S110D*); **c** Live cell imaging was used to score sister chromatid co-segregation during anaphase I in cells carrying heterozygous *CEN5-*GFP foci and Dsn1 Box 3 mutations as described in Fig. 2c, d. Data for wild type and *mam1Δ* is reproduced from Fig. 2d, other strains analyzed and number of cells counted were AMy25762 (*DSN1-V104A F107A*) n=51, AMy26475 (*DSN1-V104D F107D*) n=61 and AMy26828 (*DSN1-S109A S110A*) n=50, AMy27009 (*DSN1-S109D* S110D) n=64; **d, e** Analysis of Mam1-9Myc association with a representative centromere (*CEN4*) by anti-Myc chromatin immunoprecipitation followed by quantitative PCR (ChIP-qPCR). Wild type (AM25617), *DSN1-V104A F107A* (AM24669), *DSN1-V104D F107D* (AMy26778), *DSN1-S109A S110A* (AMy26800) and *DSN1-S109D S110D* (AMy26476) cells carrying *MAM1-9MYC* were arrested in metaphase I of meiosis by depletion of Cdc20. Strain AMy8067 was used as a no tag control. Shown is the average from 8 biological replicates for wild type and no tag. The average from 3 experiments is shown for all *DSN1* mutants with the exception of *DSN1-S109D S110D* where the average from 5 biological replicates is shown. Error bars indicate standard error; **e** Closeup view of the *Cg* Dsn1 Box 3 region with conserved serine residues mutated to aspartate (from the structure of *Cg* Csm1^69-181^:*Cg* Dsn1^43-67^DD). Residue D66 is visible forming hydrogen-bond interactions with Csm1 K172. The side-chain for residue D67 is disordered, and is modeled as alanine. Csm1 is shown in white with surface colored by charge

### Phosphorylation of Dsn1 Box 3 residues may stabilize Csm1 binding

Dsn1 Box 3 contains two serine residues (*Sc* Dsn1 S109 and S110) that are highly conserved throughout budding yeast (Fig. S6a, b). These serine residues are disordered in both crystal structures but are positioned close to conserved lysine residues in Csm1 (*Cg* Csm1 K172, K175, and K179) (Fig. 3a; Fig. S2a-c, g-h). The high conservation and physical proximity of these serine residues to positively-charged residues on Csm1 suggests that these residues may become phosphorylated, and that phosphorylation could reinforce the observed binding mode between Dsn1 Box 2-3 and Csm1. Supporting this idea, we could reconstitute a complex of *Cg* Csm1^69-181^ and a minimized *Cg* Dsn1 Box 2-3 construct (residues 43-67) with both S66 and S67 (equivalent to *Sc* Dsn1 S109 and S110) mutated to asparate to mimic phosphorylation (referred to as *Cg* Csm1^69-181^:Dsn1^43-67^DD) (Fig. S1f). We determined a 1.8 Å-resolution structure of this complex (Table S1), which closely agrees with both structures described above with the addition of a specific interaction between Dsn1 residue D66 and Csm1 K172 (Fig. 3e; S2b, e, h; S3c).

Consistent with the idea that phosphorylation of Dsn1 Box 3 promotes Csm1 binding, we found that *Sc* Dsn1^71-110^ with the phosphomimetic S109D/S110D mutation showed increased binding to *Sc* Csm1^69-190^ *in vitro* (Fig. 1e). Furthermore, Mam1 was recruited to centromeres in cells carrying the Dsn1 S109D/S110D phosphomimetic mutations, sister *CEN5-GFP* foci segregated normally to the same pole and, consistently, spore viability was comparable to that of wild type cells whether one or both copies of Dsn1 carried the mutations (Fig. 3b-d). We also analyzed a non-phosphorylatable S109A/S110A mutant and found that although kinetochore-associated Mam1 levels in a metaphase I arrest were reduced to a level comparable to that caused by the V104A/F107A and V104D/F104D mutations (Fig. 3d), the effect on spore viability and sister chromatid co-segregation was less pronounced (Fig. 3b, c). The reasons for this difference remain unclear, but we speculate that phosphorylation of S109/S110 could be dispensable for the initial recruitment of monopolin to kinetochores, but important for its maintenance into metaphase I. Although we have yet to observe Dsn1 phosphorylation on S109/S110 *in vivo*, these findings suggest that phosphorylation of Dsn1 on conserved serine residues in Box 3 could be important for stable association of monopolin with kinetochores.

### Dsn1 Box 1 is critical for meiosis

Sequence alignments of the Dsn1 N-terminal region from across fungi revealed that a conserved Dsn1 Box 2-3 region is found only in those fungi with point centromeres (Meraldi et al. 2006; Westermann et al. 2007; Gordon et al. 2011) and a *MAM1* gene (indicating the use of monopolin to co-orient sister chromatids during meiosis) (Ye et al. 2016) (Fig. S6a, c). In contrast, Dsn1 Box 1 is highly conserved throughout fungi, suggesting a possible ancestral function (Fig. S6). A recent study has shown that Dsn1 Box 1 is dispensable for binding to Csm1 as measured by yeast two-hybrid assay (Sarkar et al. 2013), and our own biochemical reconstitutions reveal that the Box 2-3 region alone binds stably to Csm1, at least when the conserved Box 3 serine residues are mutated to aspartate (Fig. S1f). These findings have suggested that Dsn1 Box 1 may not be important for monopolin complex function in point-centromere fungi including *S. cerevisiae*.

Our structure of the *Cg* Csm1^69-181^:*Cg* Dsn1^14-72^ complex shows how Dsn1 Box 2-3 binds Csm1, but also reveals a plausible Csm1 binding mode for Box 1. In the structure, Dsn1 Box 1 extends outward from the Csm1-Dsn1 Box 2-3 complex described above, and interacts with a crystallographic symmetry-related Csm1 monomer. This interface is remarkably similar to the Box 3 interface: Box 1 inserts two conserved hydrophobic residues (*Cg* Dsn1 F32 and F34, equivalent to *Sc* Dsn1 L72 and F74) into the conserved hydrophobic cavity on Csm1, and these residues are bracketed by positively- and negatively-charged residues that make specific interactions with Csm1 as in the Box 3 interface (Fig. 4a). Interestingly, Box 1 is oriented in the opposite direction across the Csm1 hydrophobic cavity than Box 3, with positively charged residues (*Cg* Dsn1 R35 and R36, equivalent to *Sc* Dsn1 K75 and R76) C-terminal to the hydrophobic residues, and negatively charged residues (*Cg* Dsn1 D29 and E30, equivalent to *Sc* Dsn1 S29 and P30) N-terminal (Fig. 4a; Fig. S4). Thus, both Dsn1 Box 1 and Box 3 bind the same surface of Csm1, through similar but distinct interaction modes. Importantly, while a Csm1 homodimer possesses two conserved hydrophobic cavities, the Dsn1 Box 1 and Box 2-3 interfaces with Csm1 are incompatible with one another, in the sense that a single Dsn1 monomer could not form both interactions with a single Csm1 dimer (Fig. S3a).

**Fig. 4.**
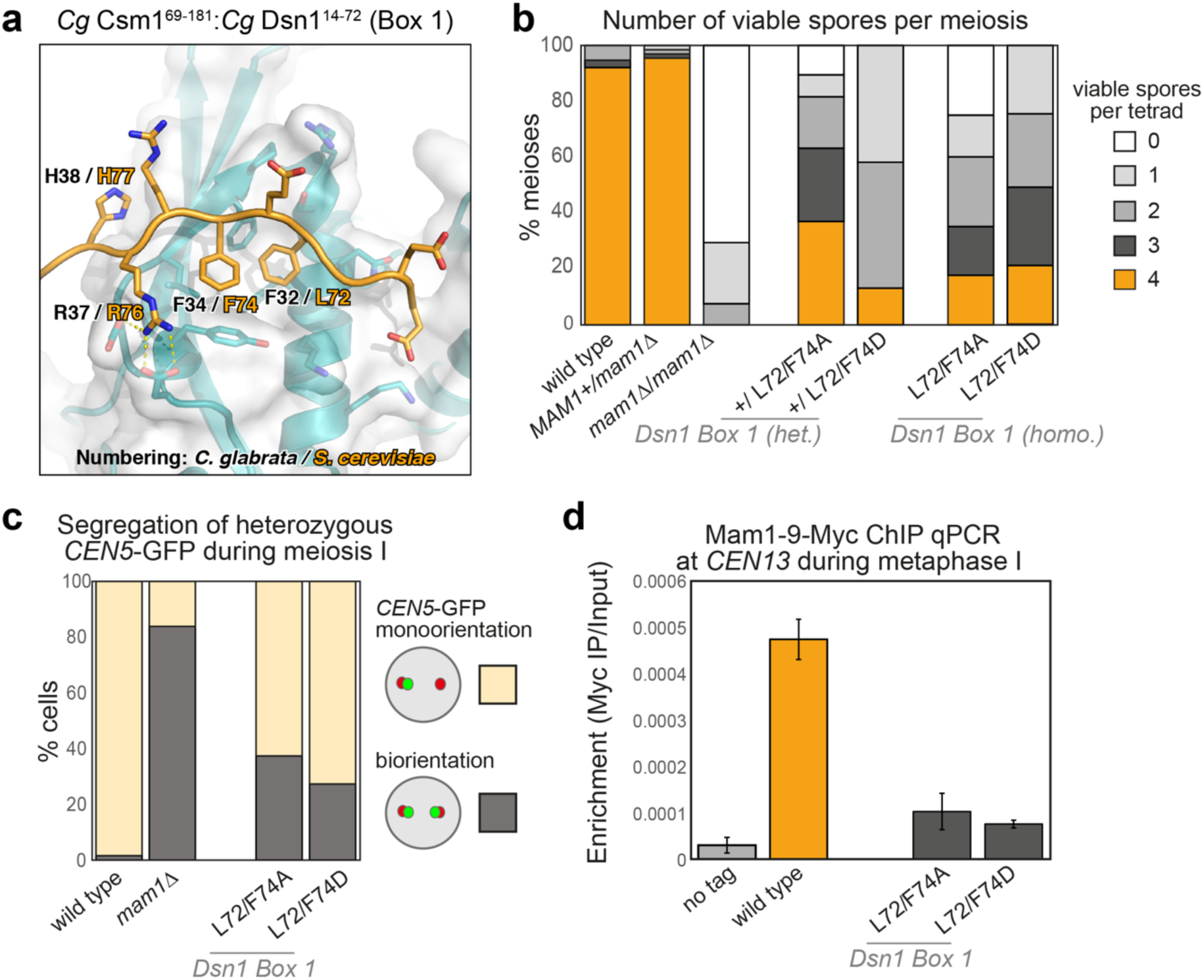
The Csm1-Dsn1 Box 1 interface. **a** Closeup view of the *Cg* Dsn1 Box 1 region (orange) interacting with the Csm1 conserved hydrophobic cavity (blue with white surface) in the *Cg* Csm1^69-181^:*Cg* Dsn1^14-72^ complex. Residue numbers shown are for *Cg* Dsn1, with *Sc* Dsn1 equivalents shown in orange text; **b** Dsn1 Box1 is critical for meiosis. Spore viability of diploid strains with the indicated genotypes were analyzed as described in Fig. 2b. Between 38 and 68 tetrads were dissected for each condition, from a minimum of two independent diploids. Data for wild type and *mam1Δ* is reproduced from Fig. 2b. Other diploids were generated from matings between AMy1827 and AMy11417 (heterozygous *mam1Δ*), AMy1827 and AMy17222 (heterozygous *DSN1-L72A F74A*), AMy1827 and AMy17123 or AMy17313 (heterozygous *DSN1-L72D L74D*), AMy17222 and AMy17223 (homozygous *DSN1-L72A F74A*) or AMy17313 and AMy17373 (homozygous *DSN1-L72D F72D*). **c** Live cell imaging was used to score sister chromatid co-segregation during anaphase I in cells carrying heterozygous *CEN5-*GFP foci and Dsn1 Box 1 mutations as described in Fig. 2c, d. Data for wild type and *mam1Δ* is reproduced from Fig. 2d, other strains analyzed were AMy25821 (*DSN1-L72A F74A*; n= 78) and AMy26543 (*DSN1-L72D F72D*; n=59); **d** Mam1 association with a representative centromere in a metaphase I arrest was analyzed by ChIP-qPCR as described in Fig 3d. Data for wild type and no tag is reproduced from Fig. 3d. Other strains used were AMy25618 (*DSN1-L72A F74A*) and AMy26543 (*DSN1-L72D F72D*) and the average of 3 or 5 biological replicates, respectively, is shown with error bars representing standard error

To test the importance of Dsn1 Box 1 for Csm1 binding and sister kinetochore monoorientation, we mutated *Sc* Dsn1 L72 and F74 to either alanine or aspartate. Both mutations reduced binding of Dsn1 to Csm1 in our *in vitro* pulldowns, with the L72D/F74D mutation having the greatest effect (Fig. 1e). Interestingly, these mutations also had a profound effect on spore viability, Mam1 association with centromeres during metaphase I, and segregation of sister chromatids during anaphase I (Fig. 4b-d). Interestingly, Dsn1 Box1 mutations caused a reduction in spore viability even when only one copy of Dsn1 carried these mutations (Fig. 4b). The presence of only a single copy of *MAM1,* in contrast, has no detectable effect on spore viability (Fig. 4b). Truncation of the first 110 amino acids of one copy Dsn1 was previously found to have a dominant-negative effect on meiosis (Sarkar et al. 2013). Our findings are consistent with this observation and further indicate that successful meiosis relies on the integrity of Box 1 in all copies of Dsn1. These findings strongly suggest that the interaction between Dsn1 Box1 and the conserved hydrophobic cavity on Csm1, as captured in the crystal structure of *Cg* Csm1^69-181^:*Cg* Dsn1^14-72^ (Fig. 4a) is important for kinetochore monoorientation *in vivo*.

### Regional-centromere fungi utilize Dsn1 Box 1 for Csm1 binding

Dsn1 Box1 is conserved in a wide range of fungi, including *S. pombe* where the monopolin subunits Csm1 and Mde4 are important for mitotic chromosome segregation, but dispensable for sister monoorientation in meiosis I (Gregan et al. 2007; Choi et al. 2009) (Fig. 5a; S6d, e). The wide conservation of Dsn1 Box 1 suggests that it may be an ancestral monopolin-binding motif that recruits the complex to the kinetochore throughout fungi, potentially through an interface like that in our structure of *Cg* Csm1^69-181^:*Cg* Dsn1^14-72^ (Fig. 5b). Supporting this idea, we have previously shown that *S. pombe* Csm1 (also called Pcs1) interacts with Dsn1 (Mis13) through its C-terminal globular domain, and that mutations to the conserved hydrophobic cavity of *S. pombe* Csm1 disrupt this interaction (Corbett et al. 2010). We used a fluorescence polarization binding assay to test whether a peptide encoding *Sp* Dsn1 Box 1 (residues 5-17) is sufficient for Csm1 binding, and measured robust binding with a *K_d_* of 22 µM (Fig. 5c). This binding was eliminated when we mutated *Sp* Csm1 isoleucine 241, located in the hydrophobic cavity, to aspartate (Fig. 5b). *Sp* Csm1 I241 is equivalent to *Sc* Csm1 L161, mutation of which we previously showed disrupts binding to Csm1 (Corbett et al. 2010). To further characterize this interaction, we used an *in vitro* pulldown assay to systematically test the importance of conserved residues in *Sp* Dsn1 Box 1 (Fig. 5d). We found that Csm1 binding was disrupted upon mutation of the central hydrophobic residues (F11 and F13), upstream negatively-charged residues (E6 and E9), and downstream positively-charged residues (R15 and K18). These data agree closely with our structure of the *Cg* Csm1:Dsn1 Box 1 interface, and strongly suggest that in *S. pombe* and likely other fungi, Dsn1 Box 1 is primarily responsible for interactions with the Csm1:Lrs4 (Pcs1:Mde4) complex.

**Fig. 5.**
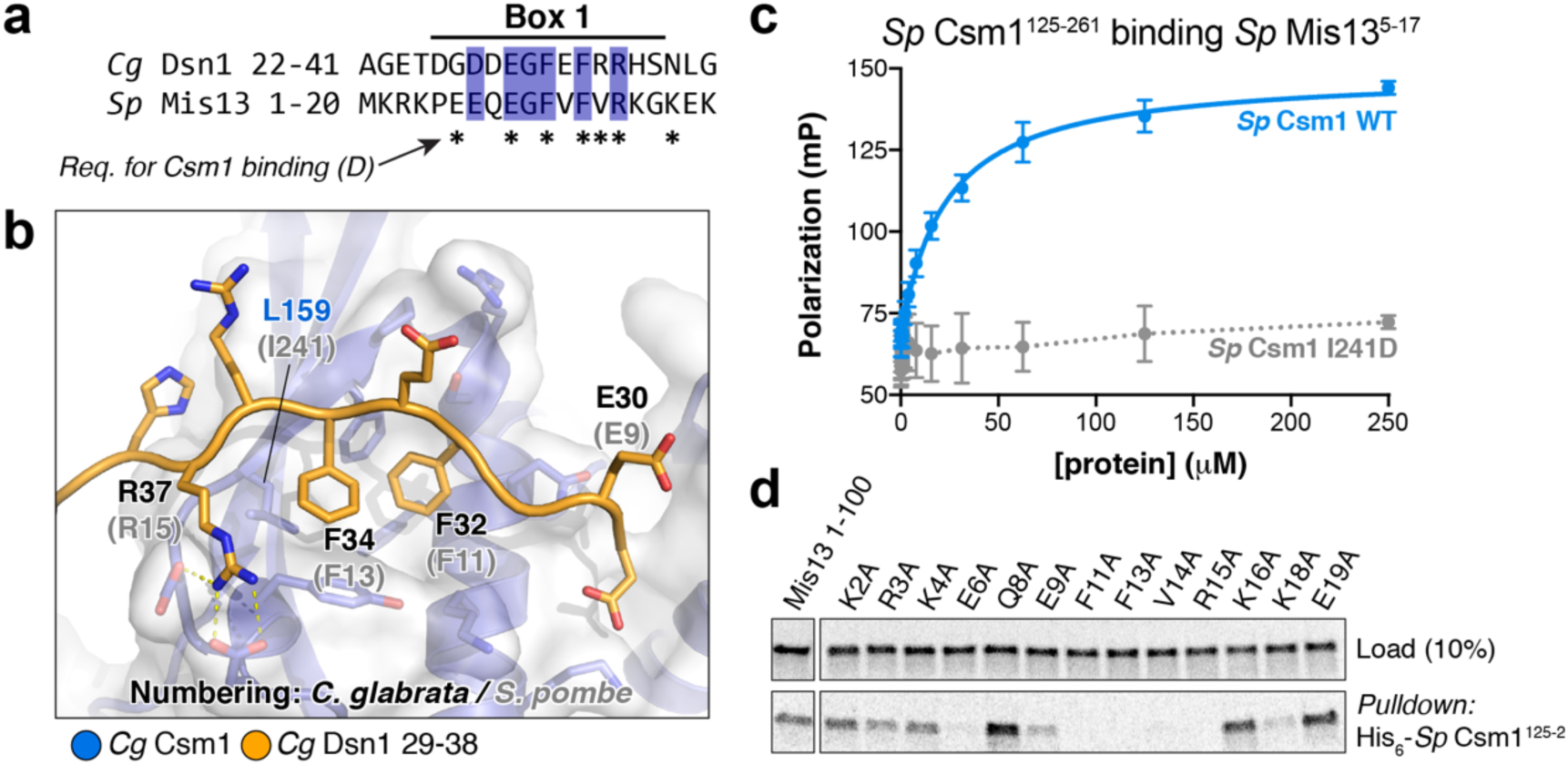
*S. pombe* Mis13 possesses a Csm1-binding Box 1 region. **a** Sequence alignment of the *Cg* Dsn1 Box 1 region with the N-terminal region of *S. pombe* Mis13. See Fig. S6 for larger sequence alignments; **b** Closeup view of the *Cg* Dsn1 Box 1 region (orange) interacting with the Csm1 conserved hydrophobic cavity (blue with white surface). Residue numbers shown are for *Cg* Dsn1 (orange) and Csm1 (blue), with *S. pombe* Mis13 equivalents shown in grey text; **c** Fluorescence polarization peptide-binding assay showing interaction of the isolated *S. pombe* Mis13 Dsn1 Box 1 region (residues 5-17, sequence PEEQEGFVFVRKG) with purified *S. pombe* Csm1 C-terminal globular domain (residues 125-261). The Csm1 I241D mutant mimics the *S. cerevisiae* L161D mutant, known to disrupt binding of Dsn1 in vitro (Corbett et al. 2010) by disrupting the conserved hydrophobic cavity (L159 in *Cg* Csm1; see panel B); **d** Ni^2+^-pulldown of in vitro translated *S. pombe* Mis13 N-terminal region (wild-type and Box 1 mutants) by His_6_-Csm1^125-261^. Mis13 residues whose mutation to alanine disrupts Csm1 binding are marked by asterisks in panel **a**

### Dsn1 Box 1 and Box 3 perform independent roles in meiosis

We found that key residues in both Box 1 and Box 3 are important for sister kinetochore monorientation and successful meiosis (Fig. 3, 4). While the full Dsn1 Box 1-2-3 region (residues 71-110) binds robustly to Csm1 *in vitro*, neither Dsn1 Box 1 (residues 71-80) or Dsn1 Box 2-3 (residues 80-110) alone bound strongly in this assay (Fig. 6a). These findings suggest that robust Csm1 binding relies on the integrity of both Box 1 and Box 3. Consistently, point mutations in either Box 1 (residues L72 F74) or Box 3 (V104 F107) strongly reduced Csm1 binding *in vitro* (Fig. 1e). However, neither set of mutations affected spore viability as severely as deletion of *MAM1* or heterozygous truncation of the first 78 (removing Box 1) or 110 (removing Box 1-2-3) residues of Dsn1 (Fig. 6b). These data raise the possibility that Dsn1 Box 1 and Box 3 could act at least partially redundantly *in vivo*. To determine whether this is the case, we combined Dsn1 mutations with alanine substitutions in the hydrophobic residues of Box 1 or Box 3 and the putative phosphorylation sites S109/S110 to generate Dsn1-L72A/L74A/V104A/F107A, Dsn1-L72A/L74A/S109A/S110A, and Dsn1 L72A/L74A/V104A/F107A/S109A/S110A. Interestingly, spore viability after meiosis where one copy of Dsn1 carries mutations in both Box 1 and Box 3 was comparable to that of strains with Dsn1 N-terminal truncations (Δ78 and Δ110; Fig. 6b). These results show that while the integrity of both Dsn1 Box 1 and Box 3 is required for robust Csm1 binding *in vitro* and functional monoorientation *in vivo*, they also have at least partially independent roles in meiosis.

**Fig. 6.**
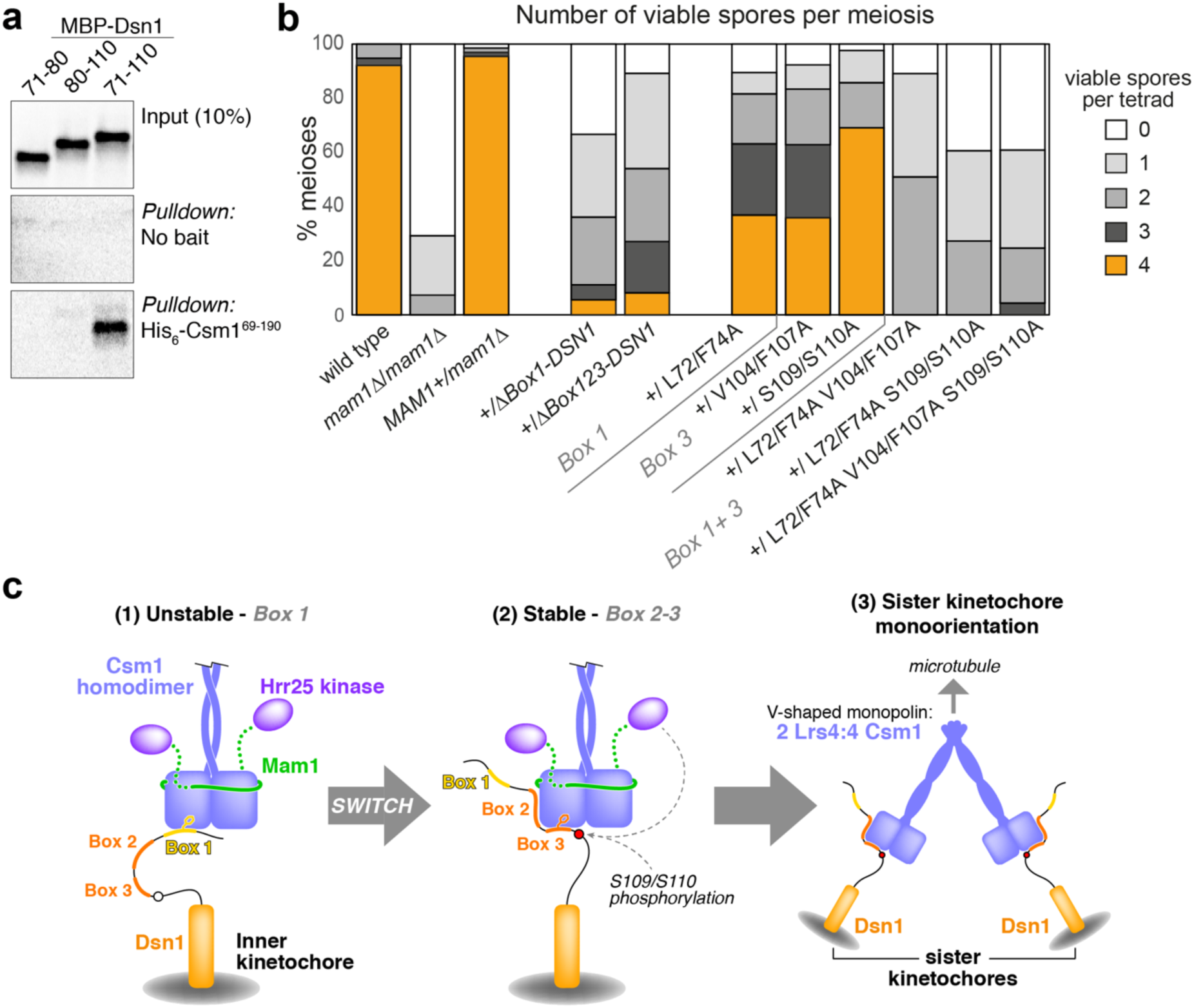
Dsn1 Box 1 and Box 3 perform independent roles in meiosis. **a** Ni^2+^-pulldown of in vitro translated *S. cerevisiae* Dsn1 N-terminal region constructs (fused to an N-terminal maltose binding protein tag) by *Sc* His_6_-Csm1^69-190^; **b** Combination of Box 1 and Box 3 mutations lead to additive effect on meiosis. Spore viability of diploid strains with the given heterozygous mutations in *DSN1* were analyzed as described in Fig. 2b. Between 38 and 68 tetrads were dissected for each Dsn1 point mutant diploid, from a minimum of two independent diploids, while more than 300 tetrads were scored for the truncations. Data for wild type and *mam1Δ* is reproduced from Fig. 2b. Other diploids were generated from matings between AMy17232 and AMy1827 (*Δ78-DSN1*), AMy17230 and AMy1827 (*Δ110-DSN1*), AMy1828 and AMy26727 (*DSN1-L72A F74A V104A F107A*), AMy1828 and AMy25883 (*DSN1-L72A F74A S109A S110A*) or AMy1828 and AMy26728 (*L72A F74A V104A F107A S109A S110A*). Data for *DSN1-L72A F74A*, *DSN1-V104A F107A*, and *DSN1-S109A S110A* is reproduced from Fig. 3b and 4c; **c** Model for sister kinetochore monoorientation by the monopolin complex. Initial unstable kinetochore association of the monopolin complex (1) occurs via Dsn1 Box 1 (yellow). While initial binding is non-specific, association of a single complex with sister kinetochores triggers a switch to a more stable binding mode (2) involving Dsn1 Box 2-3 (orange). The association is further stabilized by phosphorylation of Dsn1 S109/S110 by Hrr25 or another kinase, resulting in stable sister kinetochore monoorientation (3)

## Discussion

The V-shaped monopolin complex performs multiple roles in kinetochore and chromosome biology, which are likely to be achieved through its ability to act as a molecular crosslinker. This activity is mediated by binding of the two Csm1 homodimer heads, one at each apex of the V, to short peptide motifs on target molecules to be linked together. Our previous work had identified a conserved hydrophobic cavity on the Csm1 homodimer that is important to target monopolin both to the nucleolus and to kinetochores (Corbett et al. 2010), and a second surface important for binding additional partner proteins in both contexts (Corbett and Harrison 2012; Liang et al. 2017). Here, we defined the molecular interactions between Csm1 and its kinetochore receptor, Dsn1. We unexpectedly identify two mutually exclusive modes for Dsn1-Csm1 binding, in which the same hydrophobic cavity in Csm1 can interact with distinct conserved motifs in the Dsn1 N-terminal region (Box 1 and Box 2-3, respectively), and demonstrate that both binding modes are important for sister kinetochore monoorientation during meiosis I *in vivo*. While Box 1 is more widely conserved in fungi and likely represents the ancestral monopolin receptor at kinetochores, Box 2-3 is conserved only in yeast with point centromeres that use monopolin to direct sister kinetochore monoorientation during meiosis I.

### Dsn1 Box 1 – the ancestral monopolin recruiter

Although Mam1 orthologs appear to be present only in budding yeast with point centromeres, Csm1 and Lrs4 orthologs are found in a wide range of fungi where they are likely to function at the kinetochore in processes distinct from monoorientation during meiosis. In the fission yeast *S. pombe*, Csm1 and Mde4 associate with kinetochores during mitosis where they prevent merotelic kinetochore-microtubule attachments (Gregan et al. 2007). In budding yeast mitosis, Csm1 and Lrs4 associate with kinetochores in anaphase, independently of Mam1 (Brito et al. 2010). These observations suggest that these organisms share a conserved monopolin interaction motif in the kinetochore. Consistent with this idea, our sequence alignments revealed that Dsn1 Box 1 is broadly conserved in fungi (Fig. S6). In our structure of *Cg* Csm1^69-181^:*Cg* Dsn1^14-72^, we found that Dsn1 Box 1 forms an interaction motif that associates with the Csm1 hydrophobic cavity (Fig. 4a). Mutation of two conserved hydrophobic residues that extend into the conserved Csm1 hydrophobic cavity (L72 and F74 in *S. cerevisiae*) resulted in loss of Mam1 from kinetochores during meiosis I, a failure to properly monoorient kinetochores, and a resultant loss of spore viability (Fig. 4). This demonstrates that Dsn1 Box1 interaction with Csm1 is critical for sister kinetochore monoorientation during meiosis I. Supporting the idea that *S. pombe* monopolin uses a similar interaction mechanism for its recruitment to kinetochores, we find that the isolated Box 1 region from *Sp* Dsn1 (called Mis13) binds Csm1 via its conserved hydrophobic cavity (Fig. 5c), and that the mutation of conserved residues in *Sp* Dsn1 Box 1 disrupt binding to Csm1 (Fig. 5d). Taken together, this data provides a strong argument that the Dsn1 Box 1 represents an ancestral docking site for monopolin at kinetochores in a broad range of fungi.

### Dsn1 Box 2-3 – an adaptation to meiotic function

The Dsn1 Box 1-Csm1 interaction likely represents a general mechanism whereby monopolin can be recruited to kinetochores, but our data show that this motif on its own is insufficient for sister kinetochore monoorientation during meiosis I. Instead, both the Dsn1 Box 2-3 region and the presence of Mam1 (and by extension Hrr25) are additionally required for sister monoorientation. Structurally, Dsn1 Box 3 binds the same hydrophobic cavity on Csm1 to Dsn1 Box 1, only in the opposite orientation, with the Box 2 α-helix packing onto the side of the Csm1 homodimer. The structure of the Csm1:Dsn1 Box 3 region is also strikingly similar to that of Csm1:Tof2 (Fig. S4), which recruits Csm1 to the nucleolus to regulate ribosomal DNA silencing and recombination (Liang et al. 2017). Both Dsn1 Box 2-3 and Tof2 insert a pair of hydrophobic residues into the Csm1 hydrophobic cavity, and both possess a pair of positively-charged residues immediately N-terminal to the hydrophobic residues. Instead of two highly-conserved serine residues as in Dsn1, however, Tof2 possesses a pair of conserved negatively-charged residues that interact with the conserved lysine residues on Csm1 (Fig. S4). Thus, Csm1 interacts with diverse partners in the nucleolus and at kinetochores using similar mechanisms.

While a Csm1 homodimer possesses two identical conserved hydrophobic cavities capable of binding either Dsn1 Box 1 or Box 3, the orientation of the two binding modes and the proximity of Dsn1 Box 1 and Box 2-3 mean that it is impossible for a single Dsn1 protomer to simultaneously interact with Csm1 through both binding modes (Fig. S3a). Since the conserved Dsn1 Box 2-3 region is specifically found in budding yeast with point centromeres and Mam1 homologs, it is likely that Csm1 binding by Dsn1 Box 2-3 is specifically important for sister kinetochore monorientation. Indeed, our mutational analysis of Dsn1 Box 3 provides evidence that this is the case (Fig. 3). These observations predict the existence of meiosis I-specific mechanisms that enable Csm1-Lrs4 to engage with Dsn1 Box 2-3 and thereby fuse kinetochores. How might this be controlled? An attractive possibility is that this is the function of the Mam1-Hrr25 module of monopolin.

We previously showed that Mam1 associates with the Csm1 globular domain, and links Hrr25 to monopolin through a flexible tether (Corbett and Harrison 2012; Ye et al. 2016). In the crystal structure of *Sc* Csm1:Mam1, Mam1 forms an extended interface with Csm1 that includes an α-helix binding to the “side” of Csm1 and a phenylalanine residue (*Sc* Mam1 F262) inserted into the conserved hydrophobic cavity of Csm1 (Corbett and Harrison 2012). As these data are difficult to reconcile with our finding that Dsn1 Box 2-3 interacts with the same surfaces on Csm1, we determined a new crystal structure of a *Cg* Csm1^69-181^:Mam1^162-216^ complex (Fig. S7). This structure closely resembles our earlier structure of the *S. cerevisiae* complex, but does not include an ordered α-helix or any interaction with the Csm1 hydrophobic cavity. Closer inspection of sequence conservation in Mam1 orthologs reveals very poor conservation beyond the “core” region of Mam1 that interacts with Csm1 in our new structure, suggesting that the additional interactions observed in our earlier structure of the *S. cerevisiae* complex represented non-specific crystal packing interactions rather than a biologically relevant interface. Thus, our new structural data support the idea that Mam1 and Dsn1 Box 2-3 can bind simultaneously to a Csm1 protomer, rather than competing for Csm1 binding (Fig. S7d). Interestingly, our structural modeling suggests that Mam1 is positioned close to the solvent-exposed residues on the Dsn1 Box 2 α-helix (Fig. S7d), suggesting a potential further interaction that could stabilize the ternary complex and explain the role of these residues in Dsn1 Box 2 (Fig. 2).

Taken together, our structural and functional data suggest the following general model for monopolin function at meiosis I kinetochores (Fig. 6c): The monopolin complex may first be recruited to kinetochores through an unstable interaction between Csm1 and the ancestral Box 1 motif on the Dsn1 N-terminus. Initial binding may be non-specific, with the two Csm1 homodimer “heads” interacting with Dsn1 protomers either within a single kinetochore, between sister kinetochores, or between non-sister kinetochores. We hypothesize that sister kinetochore binding by the two Csm1 heads could trigger a switch in interaction mode between Box 1 and Box 2-3, with further stabilization of the Box 2-3 interaction coming through phosphorylation of the two conserved serine residues in Box 3 (*Sc* Dsn1 S109 and S110). Although the relevant kinase has yet to be identified, we note that Mam1 ideally positions Hrr25 to access Dsn1, and we have previously shown that Hrr25 phosphorylates Dsn1 *in vitro*, most likely in its disordered N-terminal region (Ye et al. 2016). While we did not identify specific phosphorylation sites within Dsn1, we speculate that Hrr25 may phosphorylate Dsn1 S109/S110 to stabilize monopolin-kinetochore binding. Testing the role of Hrr25, and identifying how sister kinetochore binding by the monopolin complex might be sensed, are important questions for future work.

An interesting feature of mutations in the Dsn1 Box 1-2-3 region is that they impair sister kinetochore monoorientation in a dominant fashion, despite the presence of an estimated 6-7 copies of Dsn1 at each kinetochore (Joglekar et al. 2006; Dimitrova et al. 2016). One explanation for this finding is that robust sister monoorientation relies on every copy of Dsn1 being able to associate with Csm1 through both identified binding modes, perhaps because each individual Csm1-Dsn1 interaction is relatively weak. A second possibility is that the level of Hrr25 activity at kinetochores is important, and reduced binding of monopolin in heterozygous *DSN1* mutant strains leads to a reduction in Hrr25 activity, and a consequent reduction in the efficiency of sister kinetochore monoorientation and successful meiosis.

### Monopolin – a general-purpose molecular crosslinker

While the detailed molecular mechanisms underlying specific sister kinetochore crosslinking by monopolin remain unknown, the data we present here essentially complete the structural picture of how the monopolin complex is assembled and how it interacts with kinetochores in meiosis I. When considered alongside our extensive prior structural analysis of the monopolin complex and its interactions with various binding partners, a picture emerges of a complex that is strikingly flexible in its ability to scaffold important architectural and signaling complexes in *S. cerevisiae*. The V-shaped Csm1-Lrs4 complex has two Csm1 homodimer heads positioned ~10 nm apart, and each Csm1 homodimer head possesses four protein-protein interaction surfaces: two conserved hydrophobic cavities, and two binding sites for Mam1/Ulp2-like molecules. In the nucleolus, Csm1 binds Tof2 via the conserved hydrophobic cavity, in the process likely crosslinking multiple copies of the repetitive rDNA to suppress illegitimate recombination, and aiding Sir2-dependent transcriptional silencing (Huang et al. 2006; Mekhail et al. 2008; Corbett et al. 2010; Liang et al. 2017). At the same time, Csm1 binds Ulp2 which deSUMOylates and stabilizes other key rDNA-silencing proteins (Liang et al. 2017). At the kinetochore, Csm1 binds both Dsn1 Box 1 and Box 3 via the hydrophobic cavity to directly crosslink sister kinetochores, and also binds Mam1 to indirectly recruit Hrr25. Finally, we have recently identified another Csm1 binding partner, Dse3, whose biological functions are not known but that binds Csm1 in a Ulp2/Mam1-like manner (Singh and Corbett 2018). This constellation of monopolin-interacting proteins may not yet be complete, with the *Saccharomyces Genome Database* (http://yeastgenome.org) listing 70 proteins as identified physical interactors of Csm1 alone (Singh and Corbett 2018). Thus, the monopolin complex represents a remarkably versatile molecular crosslinker that has been recruited into at least three, and potentially more, functional roles in budding yeast.

## Acknowledgements

This work was supported by Wellcome through a Senior Research Fellowship to AM [107827], core funding for the Wellcome Centre for Cell Biology [203149] and a Sir Henry Wellcome Fellowship to ED [096078]. KDC acknowledges past support from the Ludwig Institute for Cancer Research and current support from the National Institutes of Health [R01 GM104141]. RP gratefully acknowledges a studentship from the Medical Research Council. We are grateful to Weronika Borek for comments on the manuscript.

## Supplemental Information

**Table S1.**
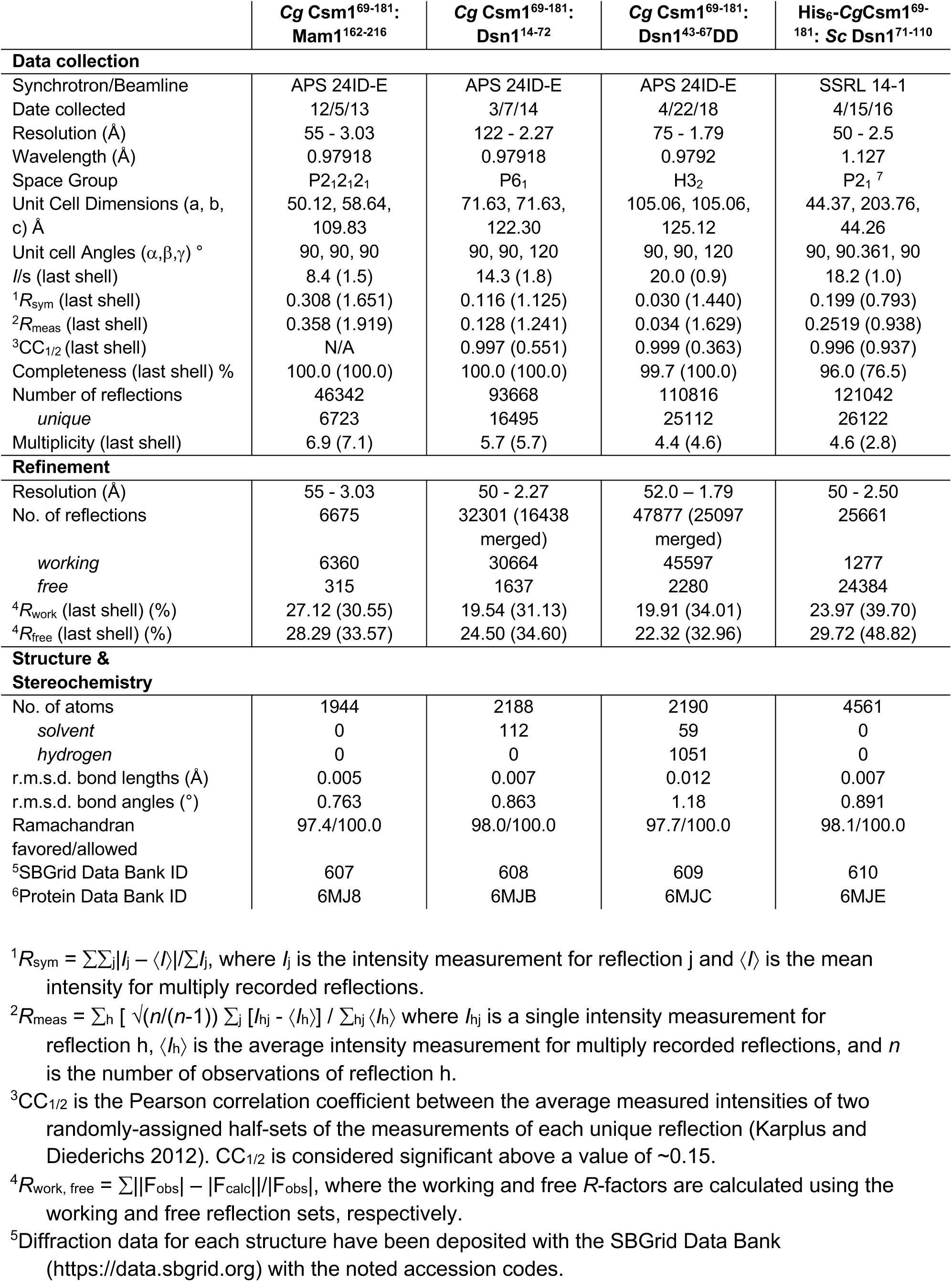
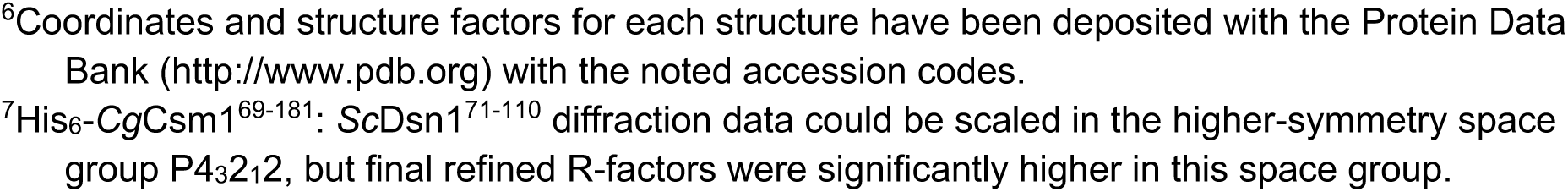
Crystallographic data collection and refinement.

|  | **Cg Csm1<sup>69-181</sup>:**<br>**Mam1<sup>162-216</sup>** | **Cg Csm1<sup>69-181</sup>:**<br>**Dsn1<sup>14-72</sup>** | **Cg Csm1<sup>69-181</sup>:**<br>**Dsn1<sup>43-67</sup>DD** | **His<sub>6</sub>-CgCsm1<sup>69-181</sup>:**<br>**Sc Dsn1<sup>71-110</sup>** |
| --- | --- | --- | --- | --- |
| **Data collection** |  |  |  |  |
| Synchrotron/Beamline | APS 24ID-E | APS 24ID-E | APS 24ID-E | SSRL 14-1 |
| Date collected | 12/5/13 | 3/7/14 | 4/22/18 | 4/15/16 |
| Resolution (Å) | 55 - 3.03 | 122 - 2.27 | 75 - 1.79 | 50 - 2.5 |
| Wavelength (Å) | 0.97918 | 0.97918 | 0.9792 | 1.127 |
| Space Group | P2<sub>1</sub>2<sub>1</sub>2<sub>1</sub> | P6<sub>1</sub> | H3<sub>2</sub> | P2<sub>1</sub><sup>7</sup> |
| Unit Cell Dimensions (a, b, c) Å | 50.12, 58.64, 109.83 | 71.63, 71.63, 122.30 | 105.06, 105.06, 125.12 | 44.37, 203.76, 44.26 |
| Unit cell Angles (α,β,γ) ° | 90, 90, 90 | 90, 90, 120 | 90, 90, 120 | 90, 90.361, 90 |
| I/s (last shell) | 8.4 (1.5) | 14.3 (1.8) | 20.0 (0.9) | 18.2 (1.0) |
| <sup>1</sup>R<sub>sym</sub> (last shell) | 0.308 (1.651) | 0.116 (1.125) | 0.030 (1.440) | 0.199 (0.793) |
| <sup>2</sup>R<sub>meas</sub> (last shell) | 0.358 (1.919) | 0.128 (1.241) | 0.034 (1.629) | 0.2519 (0.938) |
| <sup>3</sup>CC<sub>1/2</sub> (last shell) | N/A | 0.997 (0.551) | 0.999 (0.363) | 0.996 (0.937) |
| Completeness (last shell) % | 100.0 (100.0) | 100.0 (100.0) | 99.7 (100.0) | 96.0 (76.5) |
| Number of reflections | 46342 | 93668 | 110816 | 121042 |
| unique | 6723 | 16495 | 25112 | 26122 |
| Multiplicity (last shell) | 6.9 (7.1) | 5.7 (5.7) | 4.4 (4.6) | 4.6 (2.8) |
| **Refinement** |  |  |  |  |
| Resolution (Å) | 55 - 3.03 | 50 - 2.27 | 52.0 - 1.79 | 50 - 2.50 |
| No. of reflections | 6675 | 32301 (16438 merged) | 47877 (25097 merged) | 25661 |
| working | 6360 | 30664 | 45597 | 1277 |
| free | 315 | 1637 | 2280 | 24384 |
| <sup>4</sup>R<sub>work</sub> (last shell) (%) | 27.12 (30.55) | 19.54 (31.13) | 19.91 (34.01) | 23.97 (39.70) |
| <sup>4</sup>R<sub>free</sub> (last shell) (%) | 28.29 (33.57) | 24.50 (34.60) | 22.32 (32.96) | 29.72 (48.82) |
| **Structure & Stereochemistry** |  |  |  |  |
| No. of atoms | 1944 | 2188 | 2190 | 4561 |
| solvent | 0 | 112 | 59 | 0 |
| hydrogen | 0 | 0 | 1051 | 0 |
| r.m.s.d. bond lengths (Å) | 0.005 | 0.007 | 0.012 | 0.007 |
| r.m.s.d. bond angles (°) | 0.763 | 0.863 | 1.18 | 0.891 |
| Ramachandran | 97.4/100.0 | 98.0/100.0 | 97.7/100.0 | 98.1/100.0 |
| favored/allowed |  |  |  |  |
| <sup>5</sup>SBGrid Data Bank ID | 607 | 608 | 609 | 610 |
| <sup>6</sup>Protein Data Bank ID | 6MJ8 | 6MJB | 6MJC | 6MJE |

<sup>1</sup>R<sub>sym</sub> = $\sum \sum_j |I_j - \langle I \rangle| / \sum I_j$ , where $I_j$ is the intensity measurement for reflection $j$ and $\langle I \rangle$ is the mean intensity for multiply recorded reflections.
<sup>2</sup>R<sub>meas</sub> = $\sum_h [ \sqrt{n/(n-1)} \sum_j [I_{hj} - \langle I_h \rangle] / \sum_{hj} \langle I_h \rangle ]$ where $I_{hj}$ is a single intensity measurement for reflection $h$ , $\langle I_h \rangle$ is the average intensity measurement for multiply recorded reflections, and $n$ is the number of observations of reflection $h$ .
<sup>3</sup>CC<sub>1/2</sub> is the Pearson correlation coefficient between the average measured intensities of two randomly-assigned half-sets of the measurements of each unique reflection (Karplus and Diederichs 2012). CC<sub>1/2</sub> is considered significant above a value of ~0.15.
<sup>4</sup>R<sub>work, free</sub> = $\sum ||F_{obs}| - |F_{calc}|| / |F_{obs}|$ , where the working and free $R$ -factors are calculated using the working and free reflection sets, respectively.
<sup>5</sup>Diffraction data for each structure have been deposited with the SBGrid Data Bank (https://data.sbgrid.org) with the noted accession codes.
<sup>6</sup>Coordinates and structure factors for each structure have been deposited with the Protein Data Bank (http://www.pdb.org) with the noted accession codes.
<sup>7</sup>His<sub>6</sub>-CgCsm1<sup>69-181</sup>: ScDsn1<sup>71-110</sup> diffraction data could be scaled in the higher-symmetry space group P4<sub>3</sub>2<sub>1</sub>2, but final refined R-factors were significantly higher in this space group.

**Table S2.** Yeast strains used in this study.

| Strain number | Haploid/<br>Diploid | Genotype |
| --- | --- | --- |
| AMy1827 | haploid | *MATa* |
| AMy1828 | haploid | *MATalpha* |
| AMy1835 | diploid | *MATa MATalpha* |
| AMy1932 | haploid | *MATa mam1Δ::TRP1* |
| AMy1947 | haploid | *MATalpha mam1Δ::TRP1* |
| AMy8067 | haploid | *MATa cdc20::pCLB2-CDC20::KanMX6 MATalpha cdc20::pCLB2-CDC20::KanMX6* |
| AMy8291 | haploid | *MATalpha DSN1-6HIS-3FLAG::URA3 cdc20::pCLB2-3HA-CDC20::KanMX6 MATa DSN1-6HIS-3FLAG::URA3 cdc20::pCLB2-3HA-CDC20::KanMX* |
| AMy17120 | haploid | *MATalpha dsn1::DSN1(L72A F74A)-6HIS-3FLAG::URA3 cdc20::pCLB2-3HA-CDC20::KanMX6 promURA3::TetR::GFP::LEU2 tetOx224-HIS3* |
| AMy17123 | haploid | *MATalpha dsn1::DSN1(L72D F74D)-6HIS-3FLAG::URA3 cdc20::pCLB2-3HA-CDC20::KanMX6 promURA3::TetR::GFP::LEU2 tetOx224-HIS3* |
| AMy17222 | haploid | *MATalpha dsn1::DSN1(L72A F74A)-6HIS-3FLAG::URA3* |
| AMy17223 | haploid | *MATa dsn1::DSN1(L72A F74A)-6HIS-3FLAG::URA3* |
| AMy17230 | haploid | *MATa KANMX6-Δ110DSN1-6HIS-3FLAG::URA3* |
| AMy17232 | haploid | *MATalpha KANMX6-Δ78DSN1-6HIS-3FLAG::URA3* |
| AMy17313 | haploid | *MATalpha dsn1::DSN1(L72D F74D)-6HIS-3FLAG::URA3* |
| AMy17373 | haploid | *MATa dsn1::DSN1(L72D F74D)-6HIS-3FLAG::URA3* |
| AMy21920 | haploid | *MATa dsn1::DSN1(E90A N94A D97A)-6HIS-3FLAG::URA3* |
| AMy21921 | haploid | *MATa dsn1::DSN1(L88A L92A L95A)-6HIS-3FLAG::URA3* |
| AMy22719 | haploid | *MATalpha dsn1::DSN1(L88A L92A L95A)-6HIS-3FLAG::URA3* |
| AMy23151 | haploid | *MATalpha dsn1::DSN1(E90A N94A D97A)-6HIS-3FLAG::URA3* |
| AMy23152 | haploid | *MATa dsn1::DSN1(E90A N94A D97A)-6HIS-3FLAG::URA3* |
| AMy23695 | diploid | *MATalpha cdc20::pCLB2-3HA-CDC20::KanMX6 NDC10-6HA::HIS3MX6 MAM1-9MYC::TRP1 MATa cdc20::pCLB2-3HA-CDC20::KanMX6 NDC10-6HA::HIS3MX6 MAM1-9MYC::TRP1* |
| AMy24425 | diploid | *MATalpha dsn1::DSN1(L88A L92A L95A)-6HIS-3FLAG::URA3 MAM1-9MYC::TRP1 cdc20::pCLB2-3HA-CDC20::KanMX6 MATa dsn1::DSN1(L88A L92A L95A)-6HIS-3FLAG::URA3 MAM1-9MYC::TRP1 cdc20::pCLB2-3HA-CDC20::KanMX6* |
| AMy24624 | haploid | *MATa dsn1::DSN1(V104A F107A)-6HIS-3FLAG::URA3* |
| AMy24629 | haploid | *MATa dsn1::DSN1(E90K N94K D97K)-6HIS-3FLAG::URA3* |
| AMy24632 | haploid | *MATalpha dsn1::DSN1(E90K N94K D97K)-6HIS-3FLAG::URA3* |
| AMy24633 | diploid | *MATa dsn1::DSN1(S109R S110R)-6HIS-3FLAG::URA3 MAM1-9MYC::TRP1 cdc20::pCLB2-3HA-CDC20::KanMX6 MATalpha dsn1::DSN1(S109R S110R)-6HIS-3FLAG::URA3 MAM1-9MYC::TRP1 cdc20::pCLB2-3HA-CDC20::KanMX6* |
| AMy24652 | haploid | *MATalpha dsn1::DSN1(V104A F107A)-6HIS-3FLAG::URA3* |
| AMy24669 | diploid | *MATa dsn1::DSN1(V104A F107A)-6HIS-3FLAG::URA3 MAM1-9MYC::TRP1 cdc20::pCLB2-3HA-CDC20::KanMX6 MATalpha dsn1::DSN1(V104A F107A)-6HIS-3FLAG::URA3 MAM1-9MYC::TRP1 cdc20::pCLB2-3HA-CDC20::KanMX6* |
| AMy24688 | haploid | *MATa dsn1::DSN1(S109D S110D)-6HIS-3FLAG::URA3* |
| AMy24744 | haploid | *MATalpha dsn1::DSN1(S109D S110D)-6HIS-3FLAG::URA3* |
| AMy24750 | diploid | *MATa dsn1::DSN1(E90A N94A D97A)-6HIS-3FLAG::URA3 MAM1-9MYC::TRP1 cdc20::pCLB2-3HA-CDC20::KanMX6 MATalpha dsn1::DSN1(E90A N94A D97A)-6HIS-3FLAG::URA3 MAM1-9MYC::TRP1 cdc20::pCLB2-3HA-CDC20::KanMX6* |
| AMy24755 | haploid | *Mata dsn1::DSN1(V104D F107D)-6HIS-3FLAG::URA3* |
| AMy24857 | haploid | *MATalpha dsn1::DSN1(V104D F107D)-6HIS-3FLAG::URA3 cdc20::pCLB2-3HA-CDC20::KanMX6* |
| AMy24858 | haploid | *MATa dsn1::DSN1(V104D F107D)-6HIS-3FLAG::URA3* |
| AMy25111 | diploid | *MATalpha dsn1::DSN1(L72D F74D)-6HIS-3FLAG::URA3 MAM1-9MYC::TRP1 cdc20::pCLB2-3HA-CDC20::KanMX6 MATa dsn1::DSN1(L72D F74D)-6HIS-3FLAG::URA3 MAM1-9MYC::TRP1 cdc20::pCLB2-3HA-CDC20::KanMX6* |
| AMy25617 | diploid | *MATalpha MAM1-9MYC::TRP1 cdc20::pCLB2-3HA-CDC20::KanMX6 MATa MAM1-9MYC::TRP1 cdc20::pCLB2-3HA-CDC20::KanMX6* |
| AMy25618 | diploid | *MATalpha dsn1::DSN1(L72A F74A)-6HIS-3FLAG::URA3 MAM1-9MYC::TRP1 cdc20::pCLB2-3HA-CDC20::KanMX6 MATa dsn1::DSN1(L72A F74A)-6HIS-3FLAG::URA3 MAM1-9MYC::TRP1 cdc20::pCLB2-3HA-CDC20::KanMX6* |
| AMy25750 | diploid | *MATalpha dsn1::DSN1(E90A N94A D97A)-6HIS-3FLAG::URA3 PDS1-tdTomato-KITRP1 MTW1-tdTomato::NAT MATa dsn1::DSN1(E90A N94A D97A)-6HIS-3FLAG::URA3 PDS1-tdTomato-KITRP1 MTW1-tdTomato::NAT promURA3::TetR::GFP::LEU2 tetOx224-HIS3* |
| AMy25762 | diploid | *MATalpha dsn1::DSN1(V104A F107A)-6HIS-3FLAG::URA3 PDS1-tdTomato-KITRP1 MTW1-tdTomato::NAT MATa dsn1::DSN1(V104A F107A)-6HIS-3FLAG::URA3 PDS1-tdTomato-KITRP1 MTW1-tdTomato::NAT promURA3::TetR::GFP::LEU2 tetOx224-HIS3* |
| AMy25763 | diploid | *MATa dsn1::DSN1(L88A L92A L95A)-6HIS-3FLAG::URA3 MTW1-tdTomato::NAT PDS1-tdTomato-KITRP1 MATalpha dsn1::DSN1(L88A L92A L95A)-6HIS-3FLAG::URA3 PDS1-tdTomato-KITRP1 MTW1-tdTomato::NAT promURA3::TetR::GFP::LEU2 tetOx224-HIS3* |
| AMy25821 | diploid | *MATalpha dsn1::DSN1(L72A F74A)-6HIS-3FLAG::URA3 MTW1-tdTomato::NAT PDS1-tdTomato-KITRP1 MATa dsn1::DSN1(L72A F74A)-6HIS-3FLAG::URA3 PDS1-tdTomato-KITRP1 MTW1-tdTomato::NAT promURA3::TetR::GFP::LEU2 tetOx224-HIS3* |
| AMy25881 | diploid | *MATalpha dsn1::DSN1(E90K N94K D97K)-6HIS-3FLAG::URA3 PDS1-tdTomato-KITRP1 MTW1-tdTomato::NAT MATa dsn1::DSN1(E90K N94K D97K)-6HIS-3FLAG::URA3 PDS1-tdTomato-KITRP1 MTW1-tdTomato::NAT promURA3::TetR::GFP::LEU2 tetOx224-HIS3* |
| AMy25932 | diploid | *MATalpha DSN1-6HIS-3FLAG::URA3 cdc20::pCLB2-3HA-CDC20::KanMX6 MATa DSN1-6HIS-3FLAG::URA3 cdc20::pCLB2-3HA-CDC20::KanMX PDS1-tdTomato-KITRP1 MTW1-tdTomato::NAT MATa DSN1-6HIS-3FLAG::URA3 PDS1-tdTomato-KITRP1 MTW1-tdTomato::NAT promURA3::TetR::GFP::LEU2 tetOx224-HIS3* |
| AMy26337 | diploid | *MATa DSN1-6HIS-3FLAG::URA3 Ndc80-yEGFP::KanMX MTW1-tdTomato::NAT MATalpha DSN1-6HIS-3FLAG::URA3 Ndc80-yEGFP::KanMX MTW1-tdTomato::NAT* |
| AMy26426 | haploid | *MATa dsn1::DSN1(S109A S110A)-6HIS-3FLAG::URA3* |
| AMy26457 | diploid | *MATa dsn1::DSN1(L72A F74A)-6HIS-3FLAG::URA3 Ndc80-yEGFP::KanMX MTW1-tdTomato::NAT MATalpha dsn1::DSN1(L72A F74A)-6HIS-3FLAG::URA3 Ndc80-yEGFP::KanMX MTW1-tdTomato::NAT* |
| AMy26475 | diploid | *MATalpha dsn1::DSN1(V104D F107D)-6HIS-3FLAG::URA3 cdc20::pCLB2-3HA-CDC20::KanMX6 PDS1-tdTomato-KITRP1 MTW1-tdTomato::NAT MATa dsn1::DSN1(V104D F107D)-6HIS-3FLAG::URA3 PDS1-tdTomato-KITRP1 MTW1-tdTomato::NAT promURA3::TetR::GFP::LEU2 tetOx224-HIS3* |
| AMy26476 | diploid | *MATa dsn1::DSN1(S109D S110D)-6HIS-3FLAG::URA3 MAM1-9MYC::TRP1 cdc20::pCLB2-3HA-CDC20::KanMX6 MATalpha dsn1::DSN1(S109D S110D)-6HIS-3FLAG::URA3 MAM1-9MYC::TRP1 cdc20::pCLB2-3HA-CDC20::KanMX6* |
| AMy26543 | diploid | *MATalpha dsn1::DSN1(L72D F74D)-6HIS-3FLAG::URA3 MTW1-tdTomato::NAT PDS1-tdTomato-KITRP1 MATa dsn1::DSN1(L72D F74D)-6HIS-3FLAG::URA3 MTW1-tdTomato::NAT PDS1-tdTomato-KITRP1 promURA3::TetR::GFP::LEU2 tetOx224-HIS3* |
| AMy26547 | diploid | *MATa dsn1::DSN1(V104A F107A)-6HIS-3FLAG::URA3 Ndc80-yEGFP::KanMX MTW1-tdTomato::NAT MATalpha dsn1::DSN1(V104A F107A)-6HIS-3FLAG::URA3 Ndc80-yEGFP::KanMX MTW1-tdTomato::NAT* |
| AMy26778 | diploid | *MATa dsn1::DSN1(V104D F107D)-6HIS-3FLAG::URA3 cdc20::pCLB2-3HA-CDC20::KanMX6 MAM1-9MYC::TRP1 MATalpha dsn1::DSN1(V104D F107D)-6HIS-3FLAG::URA3 cdc20::pCLB2-3HA-CDC20::KanMX6 MAM1-9MYC::TRP1* |
| AMy26800 | diploid | *MATa dsn1::DSN1(S109A S110A)-6HIS-3FLAG::URA3 MAM1-9MYC::TRP1 cdc20::pCLB2-3HA-CDC20::KanMX6 MATalpha dsn1::DSN1(S109A S110A)-6HIS-3FLAG::URA3 MAM1-9MYC::TRP1 cdc20::pCLB2-3HA-CDC20::KanMX6* |
| AMy26803 | haploid | *MATalpha dsn1::DSN1(S109A S110A)-6HIS-3FLAG::URA3* |
| AMy26828 | diploid | *MATa dsn1::DSN1(S109A S110A)-6HIS-3FLAG::URA3 MTW1-tdTomato::NAT PDS1-tdTomato-KITRP1 MATalpha dsn1::DSN1(S109A S110A)-6HIS-3FLAG::URA3 MTW1-tdTomato::NAT PDS1-tdTomato-KITRP1 promURA3::TetR::GFP::LEU2 tetOx224-HIS3* |
| AMy26848 | diploid | *MATa dsn1::DSN1(S109A S110A)-6HIS-3FLAG::URA3 Ndc80-yEGFP::KanMX MTW1-tdTomato::NAT MATalpha dsn1::DSN1(S109A S110A)-6HIS-3FLAG::URA3 Ndc80-yEGFP::KanMX MTW1-tdTomato::NAT* |
| AMy27009 | diploid | *MATa dsn1::DSN1(S109D S110D)-6HIS-3FLAG::URA3 PDS1-tdTomato-KITRP1 MTW1-tdTomato::NAT MATalpha dsn1::DSN1(S109D S110D)-6HIS-3FLAG::URA3 PDS1-tdTomato-KITRP1 MTW1-tdTomato::NAT promURA3::TetR::GFP::LEU2, tetOx224-HIS3 (Tomo's centromeric)* |

**Table S3.** Plasmids used in this study.

| Plasmid # | Characteristics |
| --- | --- |
| pSB1590 (AMp770) | *DSN1-6HIS-3FLAG, URA3* |
| AMp1134 | *DSN1(L72A F74A)-6HIS-3FLAG, URA3* |
| AMp1373 | *Dsn1 (L88A L92A L95A)-6HIS-3FLAG, URA3* |
| AMp1374 | *Dsn1 (E90A N94A D97A)-6HIS-3FLAG, URA3* |
| AMp1403 | *Dsn1 (L88D L92D L95D)-6HIS-3FLAG, URA3* |
| AMp1404 | *Dsn1 (E90K N94K D97K)-6HIS-3FLAG, URA3* |
| AMp1405 | *Dsn1 (V104D F107D)-6HIS-3FLAG, URA3* |
| AMp1406 | *Dsn1 (S109D S110D)-6HIS-3FLAG, URA3* |
| AMp1484 | *Dsn1 (V104A F107A)-6HIS-3FLAG, URA3* |
| AMp1591 | *Dsn1(L72A F74A V104A F107A)-6HIS-3FLAG, URA3* |
| AMp1592 | *Dsn1(L72A F74A S109A S110A)-6HIS-3FLAG, URA3* |
| AMp1640 | *Dsn1 (S109R S110R)-6HIS-3FLAG, URA3* |
| AMp1643 | *Dsn1 (L72A F74A V104A F107A S109A S110A)-6HIS-3FLAG, URA3* |
| Amp1683 | *Dsn1 (S109A S110A)-6HIS-3FLAG, URA3* |

**Table S4.**
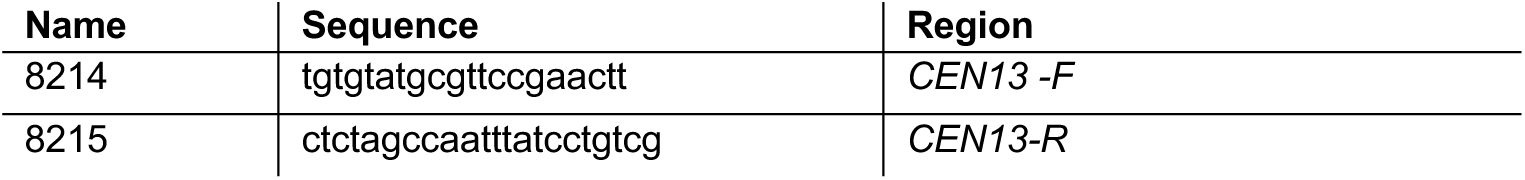
Primers used for qPCR.

**Fig. S1.**
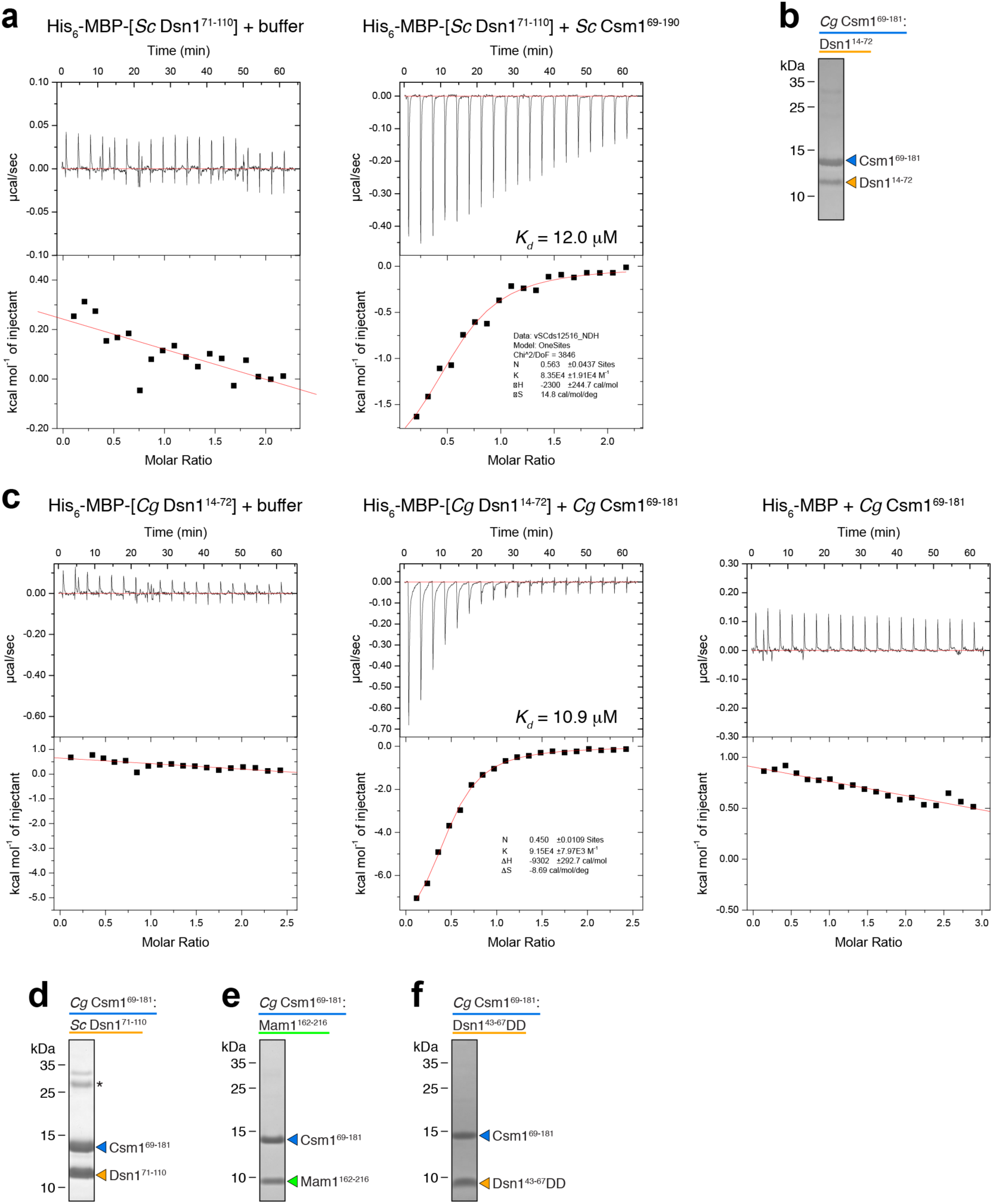
Purification and biochemical analysis of Csm1-Dsn1 complexes. **a** Isothermal titration calorimetry analysis of His_6_-MBP tagged *Sc* Dsn1^71-110^ binding *Sc* Csm1^69-190^ (*K_d_* = 12.0 µM); **b** SDS-PAGE gel of purified *Cg* Csm1^69-181^:*Cg* Dsn1^14-72^ complex; **c** Isothermal titration calorimetry analysis of His_6_-MBP tagged *Cg* Dsn1^14-72^ binding *Cg* Csm1^69-181^ (*K_d_* = 10.9 µM). *Far right:* control titration of *Cg* Csm1^69-181^ against His_6_-MBP alone. **d** SDS-PAGE gel of the *Cg* Csm1^69-181^:*Sc* Dsn1^71-110^ complex. Asterisk indicates a contaminant band. **e** SDS-PAGE gel of purified *Cg* Csm1^69-181^:*Cg* Mam1^162-216^ complex. **f** SDS-PAGE gel of purified *Cg* Csm1^69-181^:*Cg* Dsn1^43-67^DD complex

**Fig. S2.**
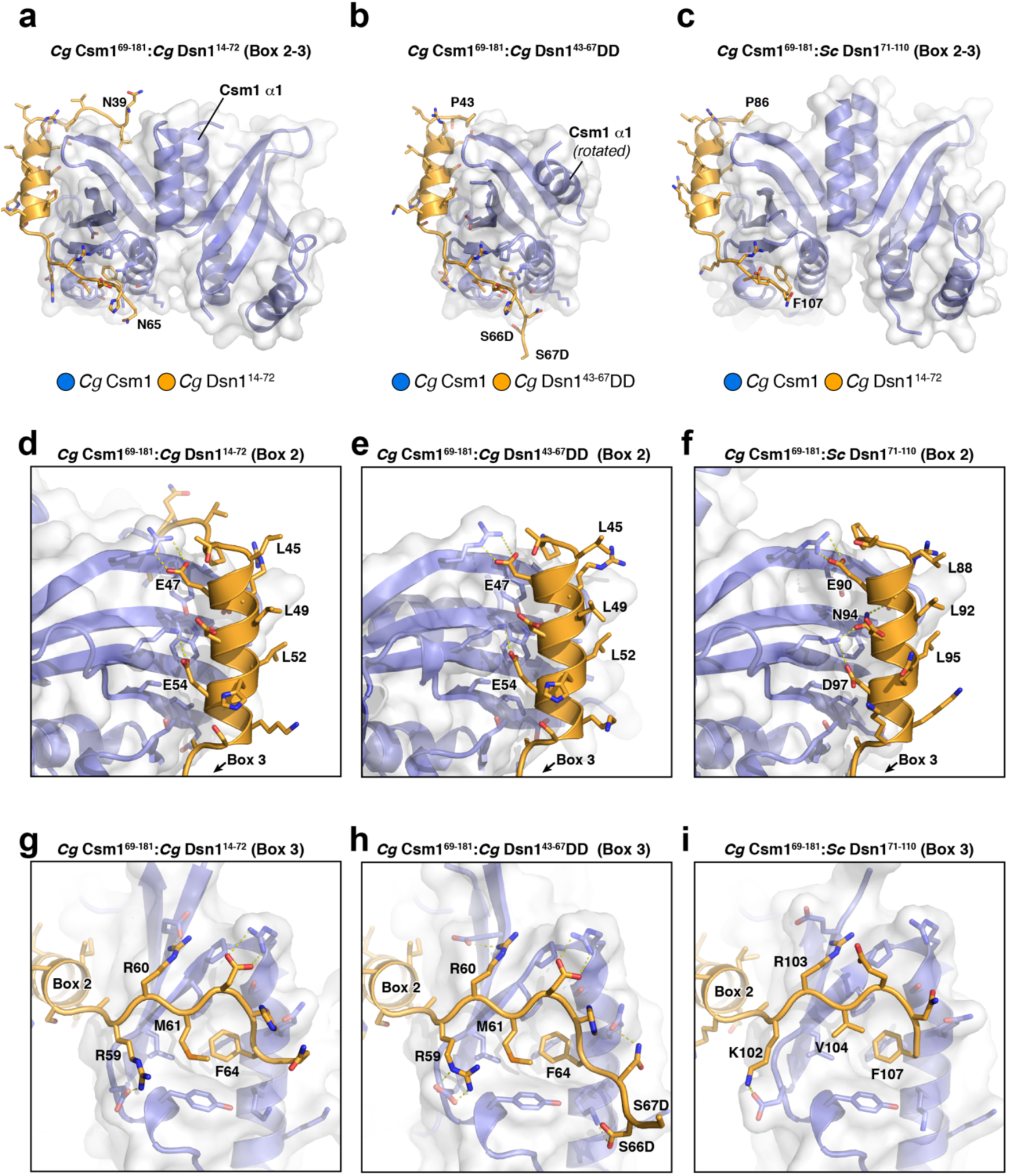
Three structures show consistent Csm1-Dsn1 Box 2-3 interactions. **a** Overall structure of the *Cg* Csm1^69-181^:*Cg* Dsn1^14-72^ complex, showing the Dsn1 Box 2-3 region; **b** Overall structure of the *Cg* Csm1^69-181^:*Cg* Dsn1^43-67^DD complex. One Csm1 protomer is shown, because of the non-canonical Csm1 dimer architecture in this crystal form (see Fig. S3c); **c** Overall structure of the *Cg* Csm1^69-181^:*Sc* Dsn1^71-110^ complex, showing the Dsn1 Box 2-3 region; **d-f** Closeup views of Dsn1 Box 2 interacting with Csm1 in *Cg* Csm1^69-181^:*Cg* Dsn1^14-72^, **d;** *Cg* Csm1^69-181^:*Cg* Dsn1^43-67^DD, **e;** and *Cg* Csm1^69-181^:*Sc* Dsn1^71-110^, **f; g-i** Closeup views of Dsn1 Box 3 interacting with Csm1 in *Cg* Csm1^69-181^:*Cg* Dsn1^14-72^, **g**; *Cg* Csm1^69-181^:*Cg* Dsn1^43-67^DD, **h**; and *Cg* Csm1^69-181^:*Sc* Dsn^171-110^, **i**

**Fig. S3.**
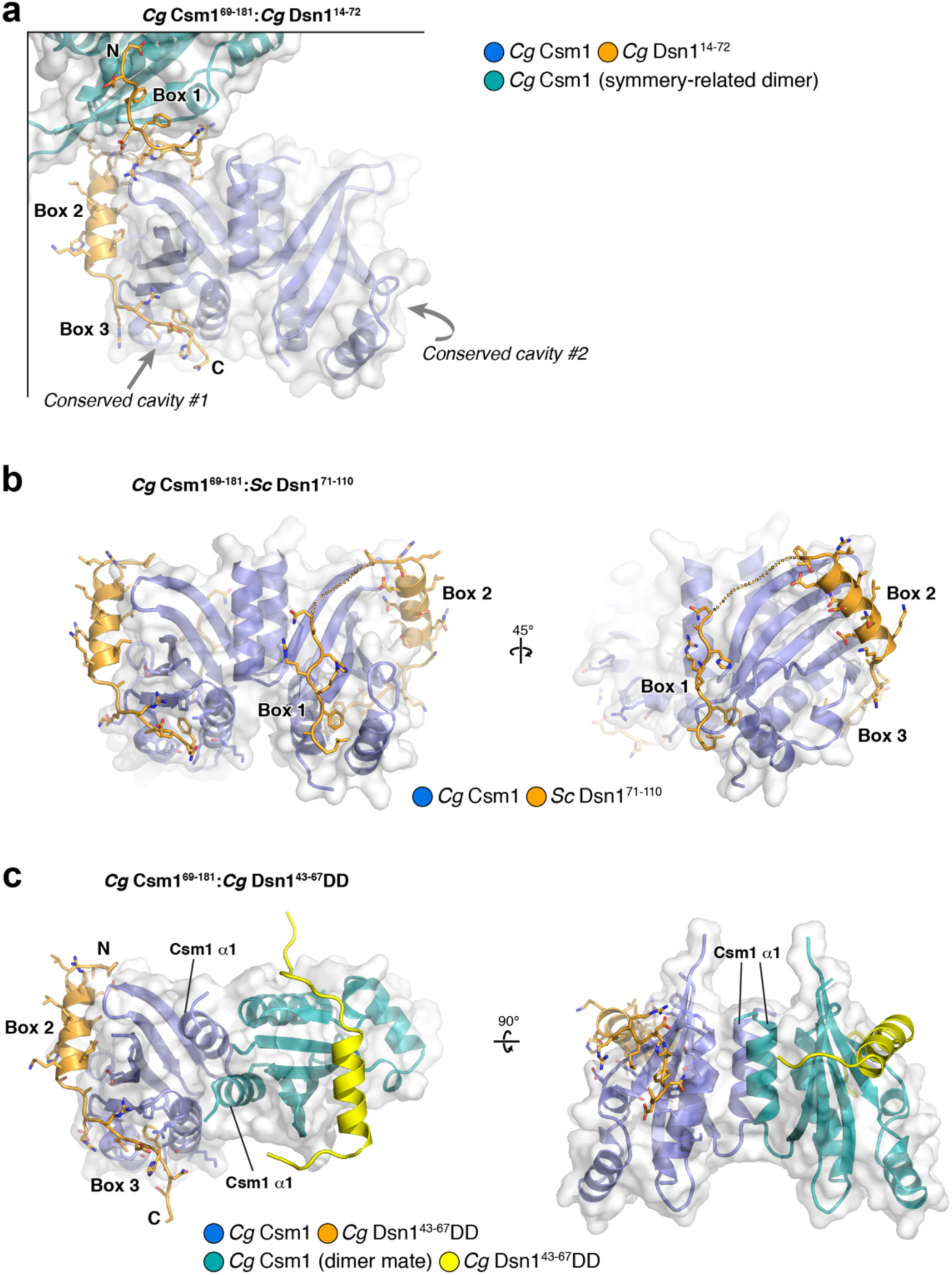
Crystal packing interactions in three structures of Csm1-Dsn1 complexes. **a** Crystal packing of the *Cg* Csm1^69-181^:*Cg* Dsn1^14-72^ complex, showing the interaction of a single Dsn1^14-72^ chain (orange with side-chains shown) with two crystallographic symmetry-related Csm1 dimers. The Dsn1 Box 2-3 region interacts with one Csm1 dimer (blue with white surface), while the Box 1 region interacts with a second crystallographic symmetry-related dimer (teal with white surface). Gray arrows indicate the locations of the two conserved hydrophobic cavities on the primary Csm1 dimer (blue). In the structure, Dsn1 Box 3 is bound to cavity #1, and cavity #2 on the dimer-related Csm1 protomer is too distant for Box 1 of the same Dsn1 protomer to simultaneously bind (this surface is occupied by Box 1 of a symmetry-related Dsn1 protomer). This crystal packing results in a 2:1 stoichiometry of Csm1 to Dsn1 in these crystals; **b** Crystal packing of the *Cg* Csm1^69-181^:*Sc* Dsn1^71-110^ complex. The asymmetric unit contains two structurally-equivalent 2:2 complexes (one shown). Each *Cg* Csm1 dimer (blue) binds two copies of *Sc* Dsn1^71-110^ (orange). In these crystals, the Dsn1 Box 1 region (residues 72-79) is ordered, and packs against a different surface of Csm1 than observed in the *Cg* Csm1^69-181^:*Cg* Dsn1^14-72^ complex. As this Csm1 surface is not well-conserved, and Dsn1 Box 1 is not tightly packed against Csm1 in this interaction (not shown), we interpret this position of Box 1 as a result of crystal packing, and not indicative of its native conformation; **b** Crystal packing of the *Cg* Csm1^69-181^:*Cg* Dsn1^43-67^DD complex. The asymmetric unit contains one Csm1 chain (blue with white surface) and one Dsn1 chain (orange with side-chains shown). The N-terminal α-helix of Csm1 is rotated ~90° from its position in all other observed Csm1 structures. A 2:2 heterotetrameric complex can be generated by crystallographic symmetry, with two Csm1 protomers interacting through their N-terminal α-lices. The unique Csm1 dimer formed by this interaction is likely the result of the low pH of the crystallization condition (pH 4.5).

**Fig. S4.**
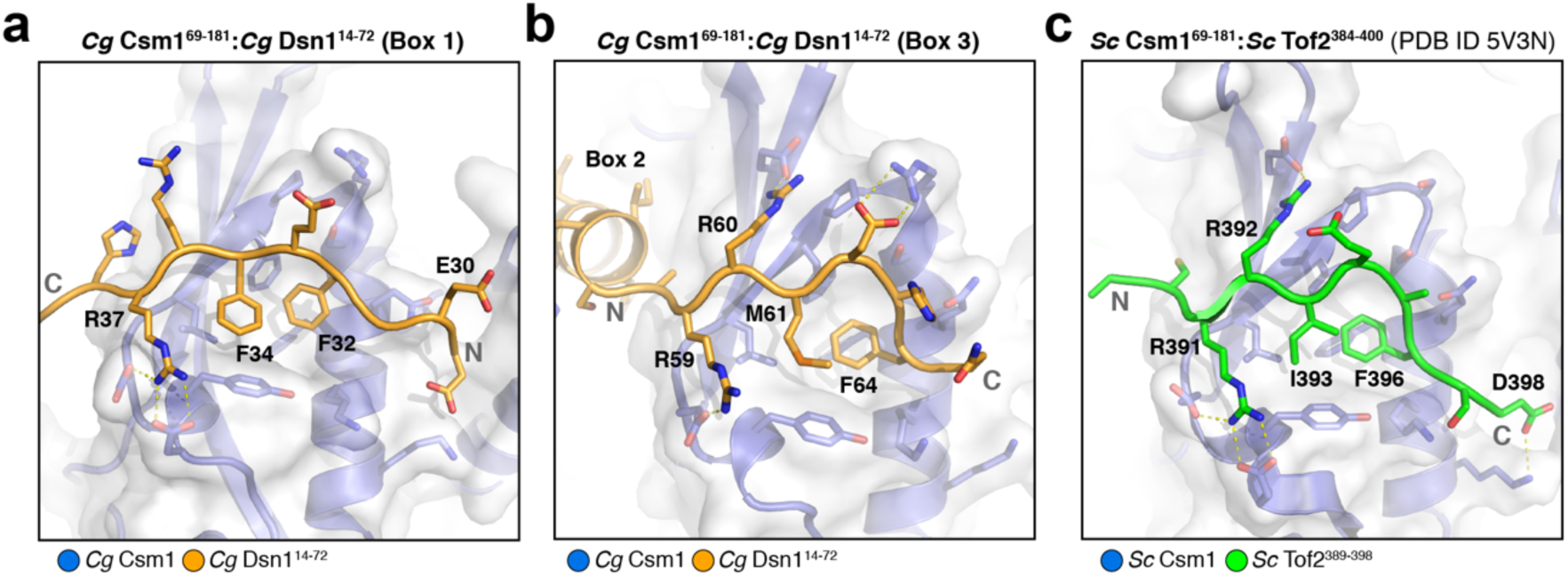
Interactions of the Csm1 conserved hydrophobic cavity with three partners. **a** Closeup view of Dsn1 Box 1 from the structure of *Cg* Csm1^69-181^:*Cg* Dsn1^14-72^, with Csm1 in blue and Dsn1 in orange; **b** Closeup view of Dsn1 Box 3 from the structure of *Cg* Csm1^69-181^:*Cg* Dsn1^14-72^, with Csm1 in blue and Dsn1 in orange; **c** Closeup view of Tof2 from the structure of Sc Csm1^69-181^:Tof2^384-400^ (PDB ID 5V3N; (Liang et al. 2017), with Csm1 in blue and Tof2 in green. Tof2 binds Csm1 equivalently to Dsn1 Box 3, while Dsn1 Box 1 is oriented in the opposite direction (N- and C-termini of each Csm1-binding motif are labeled in gray)

**Fig. S5.**
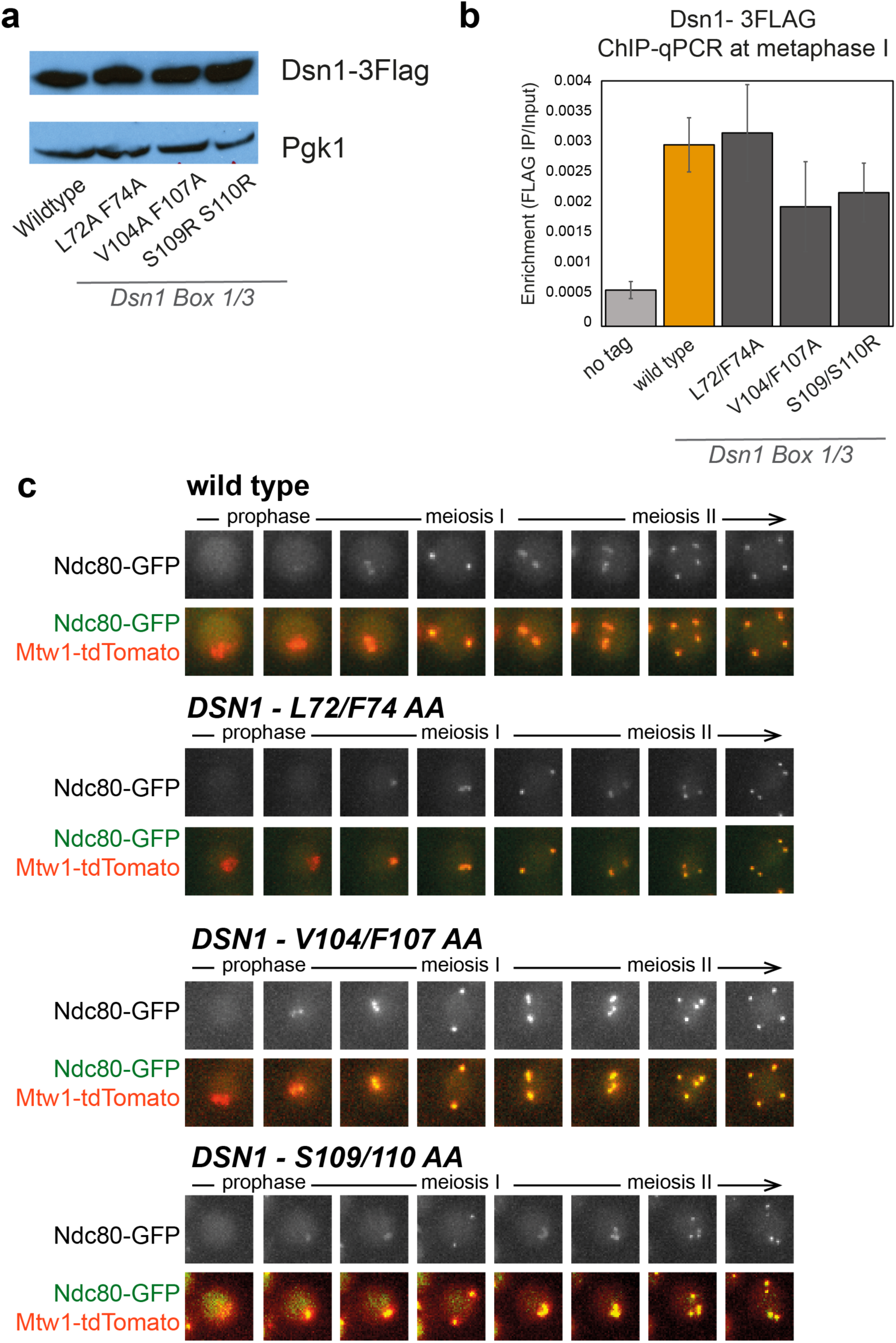
Kinetochore assembly is not affected by mutations in Dsn1’s N terminus. **a** Dsn1 mutants are expressed at wild type levels in cycling cells. Strains were as described in b and samples were isolated from exponentially growing cells; **b** Analysis of Dsn1-3Flag association with a representative centromere (*CEN13*) by anti-Flag chromatin immunoprecipitation followed by qPCR (ChIP-qPCR). Wild type (AM8291), *DSN1-L72A F74A* (Amy25618), *DSN1-V104A F107A* (Amy24669) and *DSN1-S109R S110R* (Amy24633) cells carrying *DSN1-3Flag* were arrested in metaphase I of meiosis by depletion of Cdc20. Strain AMy25617 was used as a no tag control. Shown is the average of four independent experments with error bars representing standard error. **c** Representative live cell imaging of MTW1-TdTomato to label kinetochores and Ndc80-GFP to label the outer kinetochore. Note that Ndc80-GFP is degraded in meiotic prophase and is incorporated into kinetochores only after prophase I exit (Miller et al. 2012). Images of *DSN1-L72A F74A* (AMy26457), *DSN1-V104A F107A* (AMy26547) and *DSN1-S109A S110A* (AMy26848) strains were indistinguishable from those of wild type (AMy26337), indicating that these mutations do not affect the reassembly of the outer kinetochore subunit Ndc80 after prophase exit. Images were taken at 15 minute intervals

**Fig. S6.**
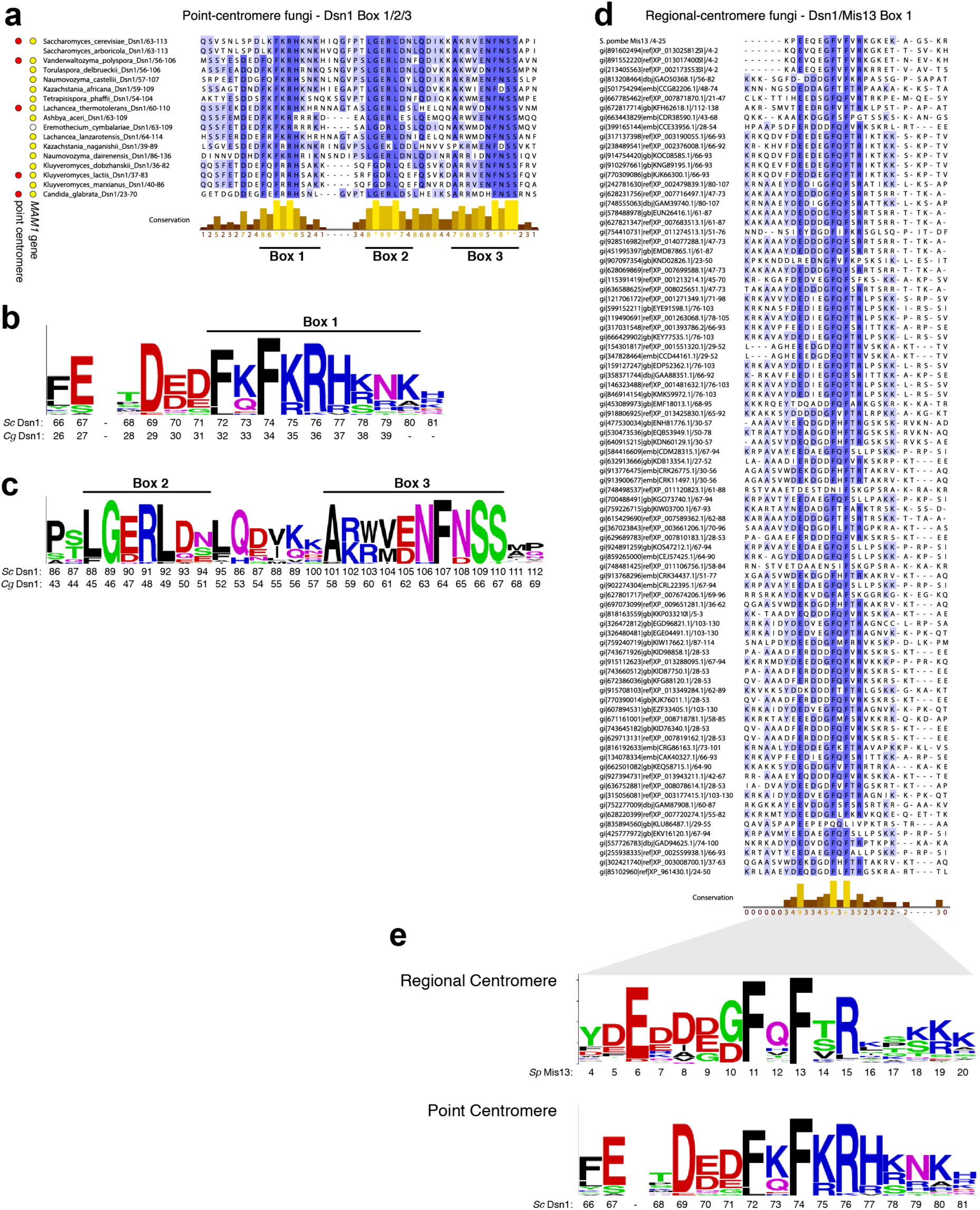
Conservation of Dsn1 Boxes 1, 2, and 3. **a** Sequence alignment of the Box 1-2-3 region of representative point-centromere fungi. Red dots on left indicate that point-centromeres have been positively identified in this organism (Meraldi et al. 2006; Gordon et al. 2011), and yellow dots indicate that the organism contains a *MAM1* ortholog (the *E. cymbalariae* contains an unannotated gene with homology to *MAM1*); **b** Sequence logo (Crooks et al. 2004) for the Dsn1 Box 1 region, generated from the alignment in panel a; **c** Sequence logo for the Dsn1 Box 2-3 region, generated from the alignment in panel a; **d** Sequence alignment of the N-terminal region of Dsn1 from 85 fungi that likely contain regional centromeres and do not possess *MAM1* orthologs; **e** *Top:* Sequence logo generated from the alignment in panel d. *Bottom:* Sequence logo for the point-centromere Dsn1 Box 1 region, from panel b, for comparison

**Fig. S7.**
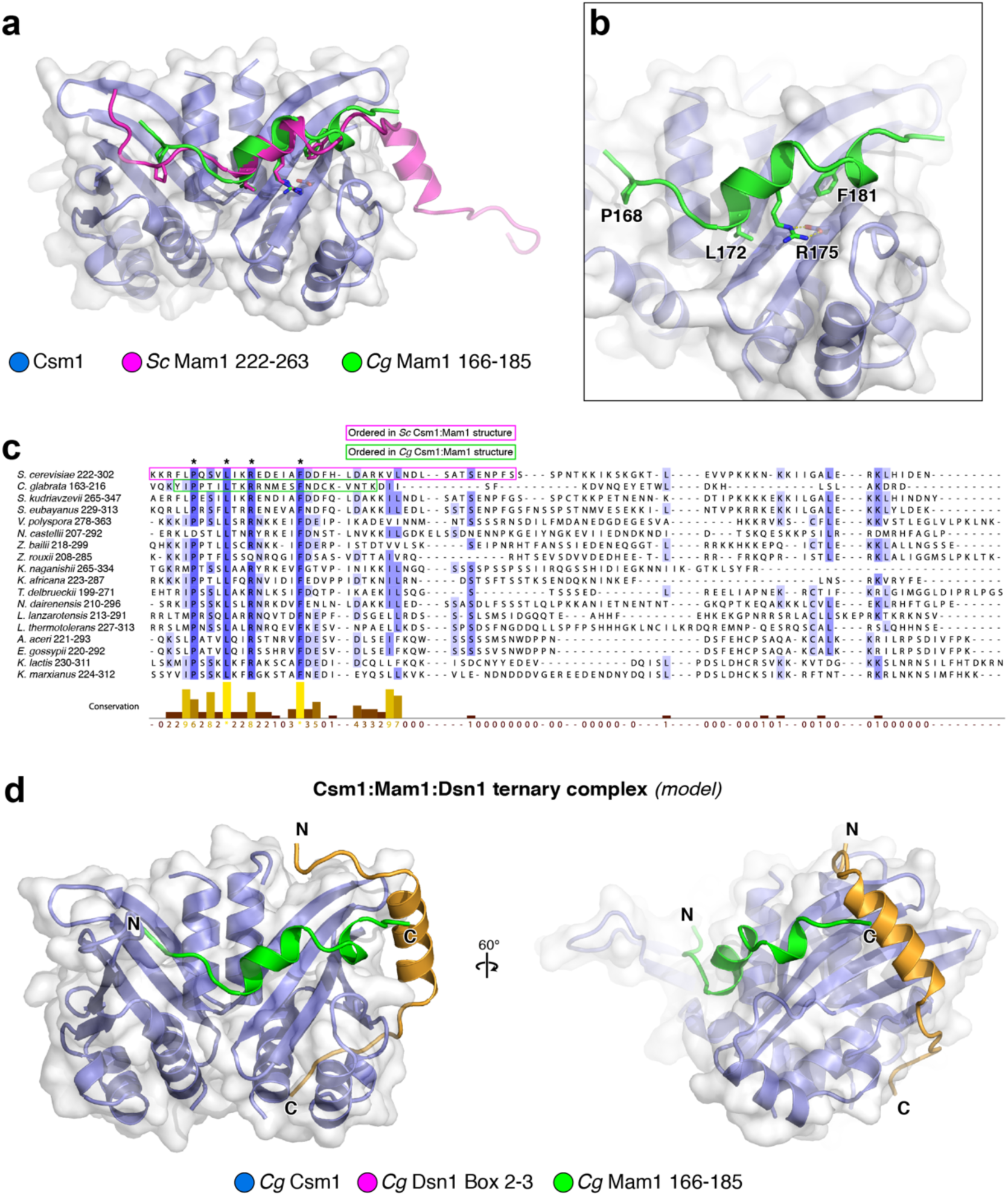
Structure of a *Cg* Csm1^69-181^:*Cg* Mam1^162-216^ complex. **a** View of the *Cg* Csm1^69-181^:*Cg* Mam1^162-216^ complex, with Csm1 shown in blue with white surface, and Dsn1 shown in green. Shown in magenta is the Mam1 chain from a prior structure of *Sc* Csm1^1-190^:*Sc* Mam1^221-302^ complex (PDB ID 5KTB) (Corbett and Harrison 2012). Shown in sticks are four highly-conserved residues in Mam1 highlighted with asterisks in panel c below; **b** Closeup view of the interaction between *Cg* Csm1 and *Cg* Mam1, with highly-conserved Mam1 residues labeled; **c** Sequence alignment of budding-yeast Mam1 C-terminal regions, highlighting the limited homology in this region and showing the ordered regions of *Sc* Mam1 (magenta) and *Cg* Mam1 (green) in their respective structures; **d** Structural model of a Csm1:Mam1:Dsn1 ternary complex, assembled by overlaying the structures of *Cg* Csm1^69-181^:*Cg* Mam1^162-216^ and *Cg* Csm1^69-181^:*Cg* Dsn1^14-72^ (Box 2-3 only). The C-terminus of Mam1 is positioned close to the Dsn1 Box 2 α-helix

